# Nonsense-mediated mRNA decay utilizes complementary mechanisms to suppress mRNA and protein accumulation

**DOI:** 10.1101/2021.05.24.445486

**Authors:** Dylan B. Udy, Robert K. Bradley

## Abstract

Nonsense-mediated mRNA decay (NMD) is an essential, highly conserved quality control pathway that detects and degrades mRNAs containing premature termination codons (PTCs). Although the essentiality of NMD is frequently ascribed to its prevention of truncated protein accumulation, the extent to which NMD actually suppresses proteins encoded by NMD-sensitive transcripts is less well-understood than NMD-mediated suppression of mRNA. Here, we describe a reporter system that permits accurate quantification of both mRNA and protein levels via stable integration of paired reporters encoding NMD-sensitive and NMD-insensitive transcripts into the AAVS1 safe harbor loci in human cells. We use this system to demonstrate that NMD suppresses proteins encoded by NMD-sensitive transcripts by up to ∼8-fold more than the mRNA itself. Our data indicate that NMD limits the accumulation of proteins encoded by NMD substrates by mechanisms beyond mRNA degradation, such that even when NMD-sensitive mRNAs escape destruction, their encoded proteins are still effectively suppressed.

## INTRODUCTION

Nonsense-mediated mRNA decay (NMD) is a eukaryotic cellular surveillance system that acts to prevent the accumulation of potentially deleterious truncated proteins by targeting mRNAs with premature termination codons (PTCs) for degradation (seminal papers: Chang et al., 1979; Losson and Lacroute, 1979; Maquat et al., 1981; Kinniburgh et al., 1982; reviewed in: Lykke-Andersen and Jensen, 2015; Kurosaki et al., 2019). In mammalian cells, mRNAs with a PTC upstream of an exon-exon junction are recognized as aberrant during translation through the interaction of the terminating ribosome with an exon junction complex (EJC) that is deposited upstream of a splice junction (Nagy and Maquat, 1998; Le Hir et al., 2000; Lykke-Andersen et al., 2001; Le Hir et al., 2001; Le Hir et al., 2016; Schlautmann and Gehring, 2020). This leads to recruitment of RNA degradation machinery that cleaves the mRNA (Huntzinger et al., 2008; Eberle et al., 2009) and thus prevents continued production of truncated proteins.

Truncated proteins derived from NMD-insensitive transcripts, in which the PTC resides in the last exon or last ∼55 nucleotides of the penultimate exon, can cause disease in heterozygotes, while heterozygous individuals bearing PTCs that generate NMD-sensitive transcripts in the same genes are often unaffected (Holbrook et al., 2004; Miller and Pearce, 2014). These genetic findings strongly support the hypothesis that limiting potentially deleterious truncated protein accumulation is essential for cell health and homeostasis and likely one of the primary selection pressures for evolution and maintenance of the NMD pathway. Despite this hypothesized importance, levels of proteins encoded by NMD-sensitive transcripts have not been quantitatively measured to the same extent as corresponding mRNA levels. Levels of NMD-sensitive mRNAs have been extensively measured and characterized (Zhang et al., 1998; Mendell et al., 2004; Tani et al., 2012; Lindeboom et al., 2016; Colombo et al., 2017; Celik et al., 2017; Kurosaki et al., 2018; Karousis et al., 2021; Kovalak et al., 2021), clearly demonstrating that NMD suppresses mRNA levels. These reduced mRNA levels imply coincident reduction of corresponding protein levels. However, in the absence of highly quantitative protein-level measurements, the extent to which protein versus mRNA alone is suppressed remains unclear.

There are multiple lines of evidence supporting the idea that proteins encoded by NMD-sensitive transcripts have the potential to accumulate to non-negligible levels: (1) NMD is a translation dependent process, so production of some potentially deleterious proteins is required to degrade the mRNA; (2) NMD does not completely deplete NMD-sensitive transcripts from cells – some remain at 20-35% levels of corresponding NMD-insensitive transcripts (Trcek et al., 2013; Hoek et al., 2019); (3) there is evidence for a subpopulation of NMD-sensitive mRNAs that are as stable as NMD-insensitive mRNAs (Tani et al., 2012; Trcek et al., 2013; Hoek et al., 2019); (4) NMD transcripts can be translated multiple times and degradation can occur after the pioneer round of translation (Kurosaki et al., 2018; Hoek et al., 2019); (5) NMD transcripts have been found to be associated with polysomes (Kim et al., 2017; Kurosaki et al., 2018); (6) NMD transcripts can be targeted for degradation even after associating with the eIF4F complex that is involved in bulk protein synthesis (Rufener and Mühlemann, 2013; Durand and Lykke-Andersen, 2013); and (7) select proteins encoded by endogenous transcripts that are predicted to be targeted by NMD can be detected (Giorgi et al., 2007).

Much of the previous work on NMD has utilized reporter systems (Daar and Maquat, 1988; Carter et al., 1996; Zhang et al., 1998; Bühler et al., 2004; Eberle et al., 2008; Kuroha et al., 2009; Kim et al., 2017; Hoek et al., 2019) that facilitate changes to the reporter sequence to test various features (e.g., PTC location, NMD-inducing features, 3’-UTR length) in a controlled manner and precisely quantify how such features affect mRNA levels. Some systems employ protein-level measurements from NMD-sensitive reporters using fluorescent proteins or luciferase (Paillusson et al., 2005; Boelz et al., 2006; Pereverzev et al., 2015; Alexandrov et al., 2017; Baird et al., 2018; Sato and Singer, 2021), but these past studies have not directly compared mRNA and protein levels. Several studies have measured both mRNA and protein levels from NMD-sensitive reporters in yeast (Muhlrad and Parker, 1999; Kuroha et al., 2009) and human cells (Inoue et al., 2004; Boelz et al., 2006; Anczuków et al., 2008; Kang et al., 2009; Kim et al., 2017; Aksit et al., 2019), although the reporters used in human cells were not optimized for precise protein-level measurements. Intriguingly, studies in yeast indicated that protein levels can be reduced to a greater degree than mRNA levels (Muhlrad and Parker, 1999; Kuroha et al., 2009). However, in the absence of quantitative and simultaneous measurements of levels of NMD-sensitive mRNAs and their encoded proteins in human cells, the extent to which mRNA and protein suppression contribute to the overall suppression of gene expression by NMD remains unclear. Overall, these past studies highlight the need to develop NMD reporters with quantitative readouts that are suitable for use in mammalian cells.

We therefore sought to develop a system to make quantitative mRNA- and protein-level measurements in human cells in order to systematically determine how NMD sensitivity influences levels of the encoded proteins relative to their parent mRNAs.

## RESULTS

### Development of an NMD reporter system for precise quantification of mRNA and protein levels

We sought to develop a reporter system based on previously validated reporters that included new, complementary features which facilitated precise measurement of both mRNA and protein levels. Such features include: (1) protein-level measurement with a high dynamic range; (2) full-length protein domains to minimize inherent instability of a truncated protein lacking any folded domains as a potentially confounding source of variability between proteins encoded by NMD-insensitive (“control”) and NMD-sensitive (“NMD(+)”) reporters; (3) internally included, NMD-insensitive control reporters to permit accurate normalization between samples; (4) straightforward measurement of mRNA and protein stability; (5) stable integration into a “safe harbor” genomic locus to eliminate the need for repeated transient transfections—which itself can reduce NMD efficiency (Gerbracht et al., 2017)—as well as remove stochastic location of genomic integration as a potentially confounding source of variability between experiments.

We employed luciferase-based reporters (based on previously published and validated reporters; Baird et al., 2018) in order to achieve high dynamic range protein-level measurements from reporter proteins with a full-length, functional domain (***Figure 1A***). Using luminescence as the readout precludes the need for western blotting and antibodies, eliminating additional potentially confounding variables. The luciferase sequences are followed by sequences that code for either full-length beta-globin (control reporters) or truncated beta-globin with a PTC at amino acid position 39 (NMD(+) reporters) (***Figure 1A***) (Zhang et al., 1998; Baird et al., 2018).

**Figure 1.**
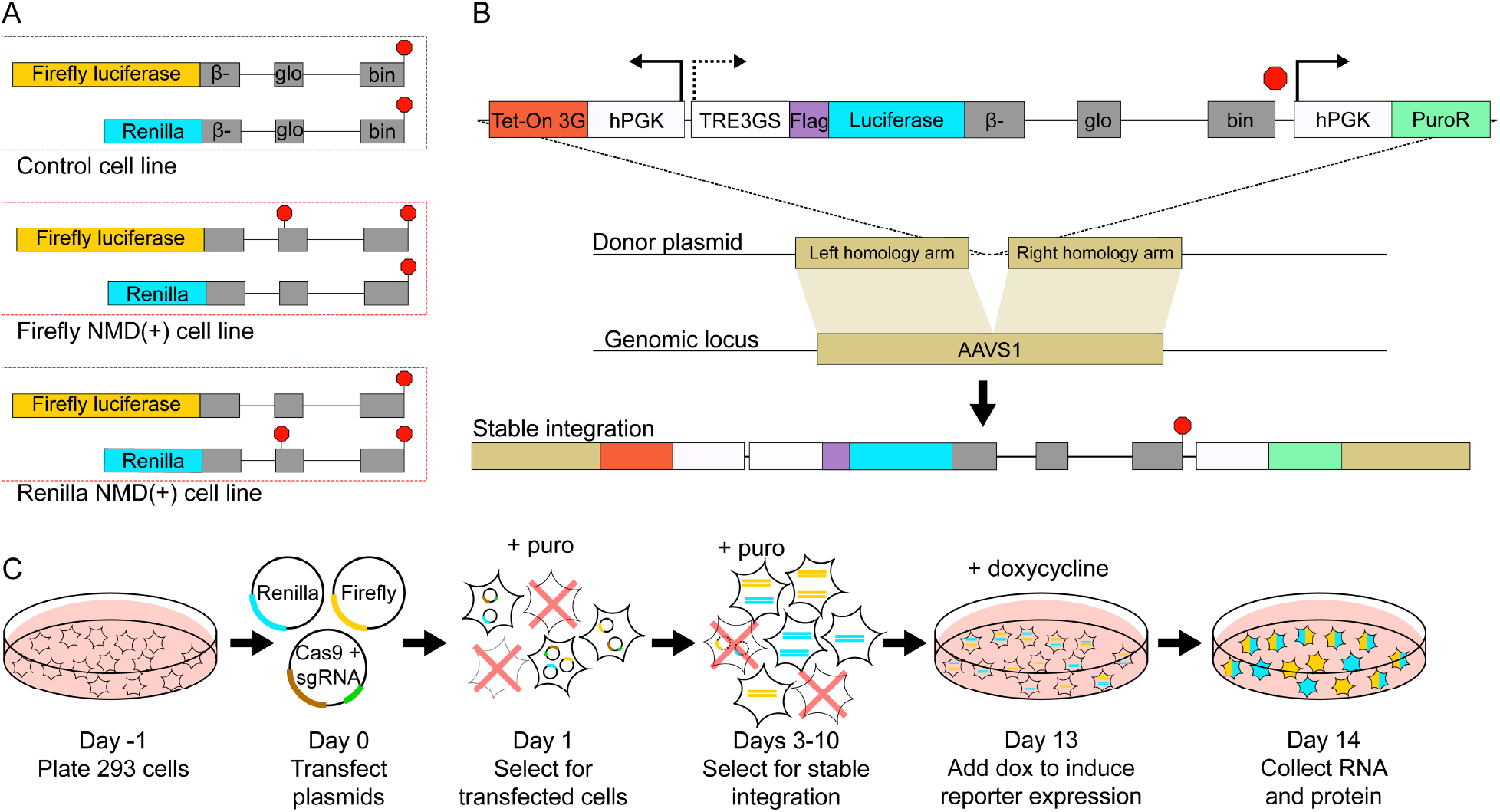
Development of a reporter system for quantitative mRNA- and protein-level measurements. (**A**) Diagrams of the luciferase-based NMD reporters used in this study. The reporters were grouped in pairs (one firefly luciferase reporter and one renilla luciferase reporter) and used together in a control cell line (both reporters have a normal termination codon) or NMD(+) cell lines (one reporter with normal TC and the other with a PTC). (**B**) Schematic of the reporter plasmid sequence that was stably integrated into the AAVS1 loci of 293 cells. (**C**) Workflow describing how the NMD reporters were stably integrated into 293 cells using CRISPR-Cas9 genome engineering and how selection for only cells with stably integrated reporters was performed.

We took advantage of the reporters’ potentiality for use in a dual-luciferase system (Sherf et al., 1996) in which two distinct luciferase enzymes (firefly and renilla) are co-expressed and one is designated as an internal control (***Figure 1A***), permitting normalization between samples with the same internal control luciferase. For example, the firefly NMD(+) reporter is normalized to a renilla control reporter in the same sample, and that ratio is then compared to the firefly control reporter normalized to the renilla control reporter in another sample to determine the firefly NMD(+) reporter level relative to the firefly control reporter level (***Figure 1—figure supplement 1***). We created two distinct NMD(+) cell lines, in which either firefly or renilla luciferase is used in the NMD(+) reporter while the other luciferase is used in the control reporter, and vice versa (***Figure 1A***, bottom two sets of reporters). This strategy ensured that results were dependent on the NMD sensitivity of the reporter rather than specific to a particular luciferase.

RNA stability is often measured using actinomycin D to inhibit transcription, which can lead to widespread changes in the transcriptome and pleiotropic effects on cell function (Lugowski et al., 2018). We therefore used a Tet-On inducible promoter system (*Gossen et al., 1995; Heinz et al., 2011*) with our NMD reporters (***Figure 1B***) to modulate reporter expression with doxycycline, enabling temporal control of expression and mRNA stability measurements without bulk transcription inhibition.

Finally, mRNAs transcribed from transiently transfected reporters are not efficiently degraded by NMD in some cell types (Gerbracht et al., 2017). To avoid such a disruptive complication and obtain more uniform and consistent reporter expression, we used CRISPR/Cas9-mediated genome engineering to stably integrate the reporters into the AAVS1 safe harbor loci in HEK-293 cells. The reporter sequences were cloned into a donor plasmid with homology arms to the AAVS1 locus (Natsume et al., 2016) (***Figure 1B***), and the donor plasmids were co-transfected with a Cas9/AAVS1-sgRNA expressing plasmid into HEK-293 cells. Cells with stably integrated reporters were selected for using puromycin over several days (workflow in ***Figure 1C***). After generation of these stable cell lines, we used RT-PCR to confirm that these reporters were efficiently and correctly spliced (***Figure 1—figure supplement 2***). We performed all subsequent experiments with these cell lines unless described otherwise.

### mRNA levels and decay kinetics confirm NMD sensitivity of the reporters

We first validated our reporters by confirming that they were subject to RNA degradation by the NMD machinery. We measured reporter mRNA levels via qRT-PCR and found that the NMD(+) reporter mRNA levels were reduced to ∼15-25% of the corresponding control reporter mRNA levels (***Figure 2A***, “siCtrl” solid boxes), a reduction similar to that observed in previous experiments that used beta-globin reporters with a PTC at amino acid position 39 (Zhang et al., 1998).

**Figure 2.**
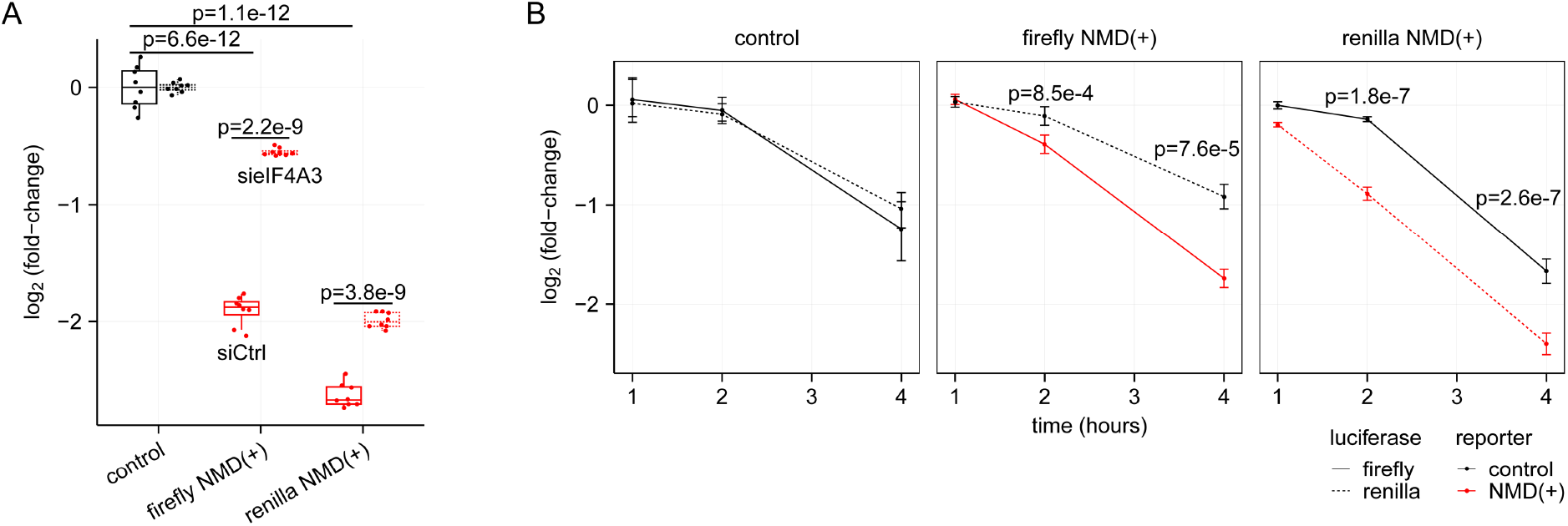
NMD reporter steady-state mRNA levels and decay kinetics are comparable to those of previously published NMD-sensitive reporters. (**A**) Box plots showing the steady-state NMD reporter mRNA levels relative to the levels in the control cell lines with and without NMD inhibition via RNAi-mediated eIF4A3 depletion. Each box plot shows n=4 technical replicates from n=2 biological replicates for a total of eight data points. An unpaired two-samples t-test was used for calculating the p-values. (**B**) Line plots showing the decay kinetics of the NMD reporter mRNA after doxycycline removal to turn off reporter transcription. The firefly and renilla reporters are plotted as separate lines. The levels at each time point are plotted relative to the levels at time point 0. The “control” panel is a combination of two independent control cell lines (both cell lines have the same two control reporters integrated, but the lines were generated separately with CRISPR/Cas9 mediated genome engineering). Each time point corresponds to n=4 technical replicates (n=2 biological replicates for the control panel, n=8 data points), with error bars showing the range of those values and the line plot connecting at the mean of the values. An unpaired two-samples t-test was used for calculating the p-values, which used the ratio of individual firefly replicate values to the mean renilla value at each time point for the NMD(+) reporter cell lines compared to the ratios at the same time point in the control cell lines. The data is plotted starting at one hour after doxycycline removal because there is little change in the reporter levels between the zero- and one-hour time points, likely due to technical limitations of the dox-inducible promoter.

To confirm that the lower mRNA levels of the NMD(+) reporters are a consequence of the desired NMD sensitivity of the transcript, we inhibited NMD by depleting eIF4A3 (***Figure 2— figure supplement 1***). eIF4A3 is a core component of the exon junction complex (EJC) (Chan et al., 2004; Palacios et al., 2004; Shibuya et al., 2004; Ferraiuolo et al., 2004) and binds directly to both spliced RNA and other core EJC factors (Shibuya et al., 2004; Bono et al., 2006; Andersen et al., 2006). Depletion of eIF4A3 is predicted to reduce EJC deposition on spliced RNAs and leads to preferential stabilization of NMD-sensitive transcripts (Palacios et al., 2004; Shibuya et al., 2004; Ferraiuolo et al., 2004; Giorgi et al., 2007). NMD(+) reporter mRNA levels increased with eIF4A3 depletion (***Figure 2A***, “sieIF4A3” dashed boxes) by up to ∼3-fold relative to control siRNA samples, confirming that reduced steady-state levels arise from action of the NMD machinery.

To determine if faster RNA degradation was responsible for the observed lower steady-state NMD(+) reporter mRNA levels, we turned off transcription using the inducible promoter to directly measure reporter mRNA decay kinetics. The NMD(+) reporter mRNA was degraded faster than was the control reporter mRNA (***Figure 2B***), as expected and consistent with previous studies (Trcek et al., 2013; Kim et al., 2017; Askit et al., 2019). We observed faster degradation for both the firefly and renilla NMD(+) reporters (***Figure 2B***, right two panels), although there were modest differences in the magnitudes of the changes. We observed these differences in magnitude for both the steady-state mRNA levels and mRNA degradation rates (***Figure 2A-B***), suggesting that they may arise from the different luciferase CDSs used in each reporter. These CDS-specific differences highlight the importance of controlling for CDS identity when studying NMD, a control that is inherent to our reporter system given its use of CDS-matched NMD-sensitive and NMD-insensitive transcripts.

Overall, these data confirm that our stably integrated reporters are modulated by NMD at the RNA level and that NMD activity suppresses their steady-state levels and influences their decay kinetics as expected based on results from previously published NMD reporters.

### NMD(+) reporter protein levels are reduced to a greater degree than are mRNA levels

We next took advantage of the reporters’ luminescence to make precise and quantitative measurements of protein levels. We inhibited NMD with RNAi of multiple NMD factors and qualitatively assessed changes in firefly NMD(+) reporter protein levels via western blot (***Figure 2—figure supplement 1***). In control siRNA samples, we observed a very faint band corresponding to the firefly luciferase plus truncated beta-globin fusion protein (***Figure 2— figure supplement 1***, lanes 1-2). The band intensity increased to the greatest degree with depletion of eIF4A3 (***Figure 2—figure supplement 1***, lanes 5-6).

For more precise quantification of these changes, we used the dual-luciferase assay to measure protein levels in the NMD reporter cell lines and normalized to control siRNA conditions and the control cell line. The protein levels for both NMD(+) reporters increased ∼6-8-fold with depletion of eIF4A3 (***Figure 3A***). Surprisingly, the NMD(+) reporter protein levels did not increase substantially with depletion of either UPF1 or SMG1 (***Figure 3A***). We plotted the data from the control and eIF4A3 siRNA conditions and normalized to just the control cell line to quantify the steady-state protein levels (***Figure 3B***). As expected, NMD(+) reporter protein levels were lower than control reporter levels under control siRNA conditions (***Figure 3B***, green solid boxes compared to black solid box).

**Figure 3.**
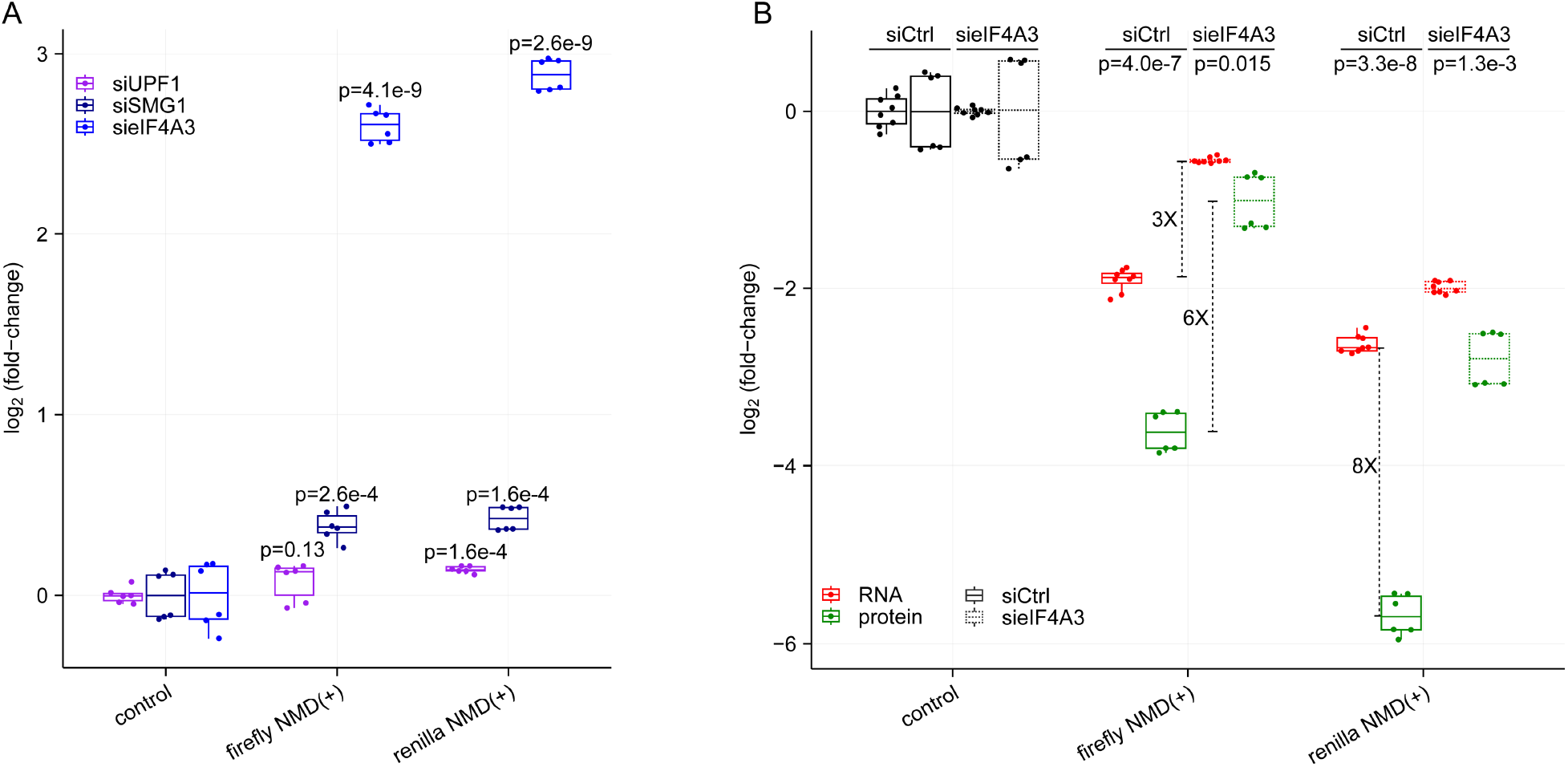
NMD(+) reporter protein levels are reduced relative to control reporter protein levels and to a greater degree than NMD(+) reporter mRNA levels. (**A**) Box plots showing the increase in NMD(+) reporter protein levels relative to control reporter protein levels upon depletion of NMD factors UPF1, SMG1, and eIF4A3. The dual-luciferase assay was used to measure reporter protein levels. Each box plot shows n=3 technical replicates normalized to n=2 biological replicates for a total of six data points. An unpaired two-samples t-test was used for calculating the p-values, which correspond to the comparison between the control cell line and each NMD(+) reporter cell line for each siRNA. (**B**) Box plots showing the comparison between NMD(+) reporter mRNA and protein levels relative to control reporter levels, with and without eIF4A3 depletion. Fold-changes are shown for the difference between mRNA and protein levels under control conditions (∼8-fold), the difference between mRNA levels with and without eIF4A3 depletion (∼3-fold), and the difference between protein levels with and without eIF4A3 depletion (∼6-fold). An unpaired two-samples t-test was used for calculating the p-values, which correspond to the comparison between mRNA and protein levels under the same siRNA treatment conditions for each NMD(+) reporter cell line.

Although decreased NMD(+) reporter protein levels relative to control reporter levels were expected, the dramatic extent of this protein-level suppression was surprising. We therefore tested whether differential rates of integration of the reporters was responsible for these pronounced differences. We assessed whether this phenomenon was still observed in a more controlled genetic setting in which exactly one copy of both the control and NMD(+) reporter was stably integrated in every cell. We performed single-cell sorting, established monoclonal cell lines, and selected clones which we confirmed via gDNA PCR had both firefly and renilla luciferase reporters stably integrated at the loci of the two AAVS1 alleles (***Figure 3—figure supplement 1***, additional details in Materials and methods). Monoclonal cell line protein levels mimicked those observed in the polyclonal lines (***Figure 3—figure supplement 2***), confirming that biased integration was not the source of the marked protein-level suppression.

We next utilized the quantitative nature of our reporters to compare the relative suppression of mRNA and protein as a consequence of NMD. Unexpectedly, the NMD(+) reporter protein levels were consistently reduced to a greater degree than were the NMD(+) reporter mRNA levels (***Figure 3B***, green solid boxes compared to red solid boxes) relative to control reporters. For example, the renilla NMD(+) reporter mRNA was reduced to ∼15% of control reporter mRNA levels, while the renilla NMD(+) reporter protein was reduced to ∼2% of control reporter protein levels (∼8-fold difference between mRNA- and protein-level suppression, annotated in ***Figure 3B)***.

We next tested whether this enhanced protein-level suppression arose from the NMD sensitivity of the reporter mRNA. We depleted eIF4A3 to inhibit NMD and measured protein levels. Upon eIF4A3 depletion, NMD(+) reporter protein levels increased relative to control siRNA conditions (***Figure 3B***, green dashed boxes vs green solid boxes). NMD(+) reporter protein levels increased to a greater degree than did NMD(+) reporter mRNA levels under eIF4A3 depletion conditions (∼6-fold protein-level increase vs ∼3-fold mRNA-level increase following eIF4A3 depletion, annotated in ***Figure 3B***). Furthermore, the NMD(+) reporter protein levels approached the same level as the NMD(+) reporter mRNA levels under eIF4A3 depletion conditions (green dashed boxes vs red dashed boxes), which is in stark contrast to the large differences under control siRNA conditions (green solid boxes vs red solid boxes). These data imply that the pronounced difference in relative suppression of mRNA and protein levels is dependent on reporter NMD sensitivity.

### NMD(+) reporter proteins are degraded modestly faster than are control reporter proteins

The greater degree of NMD(+) reporter protein reduction relative to mRNA reduction implies the existence of cellular mechanisms beyond RNA decay that limit the levels of proteins translated from NMD-sensitive transcripts. We therefore sought to test possible mechanisms responsible for this phenomenon.

One potential mechanism is through increased degradation of proteins encoded by NMD-sensitive mRNAs, which has been observed for truncated proteins encoded by NMD-sensitive reporter transcripts in yeast (Kuroha et al., 2009). To directly measure the decay kinetics of NMD reporter proteins, we inhibited translation with cycloheximide in our NMD reporter cell lines and measured protein levels at several later time points. Over the full six-hour time course there was minimal change between control and NMD(+) protein levels (***Figure 4A***).

**Figure 4.**
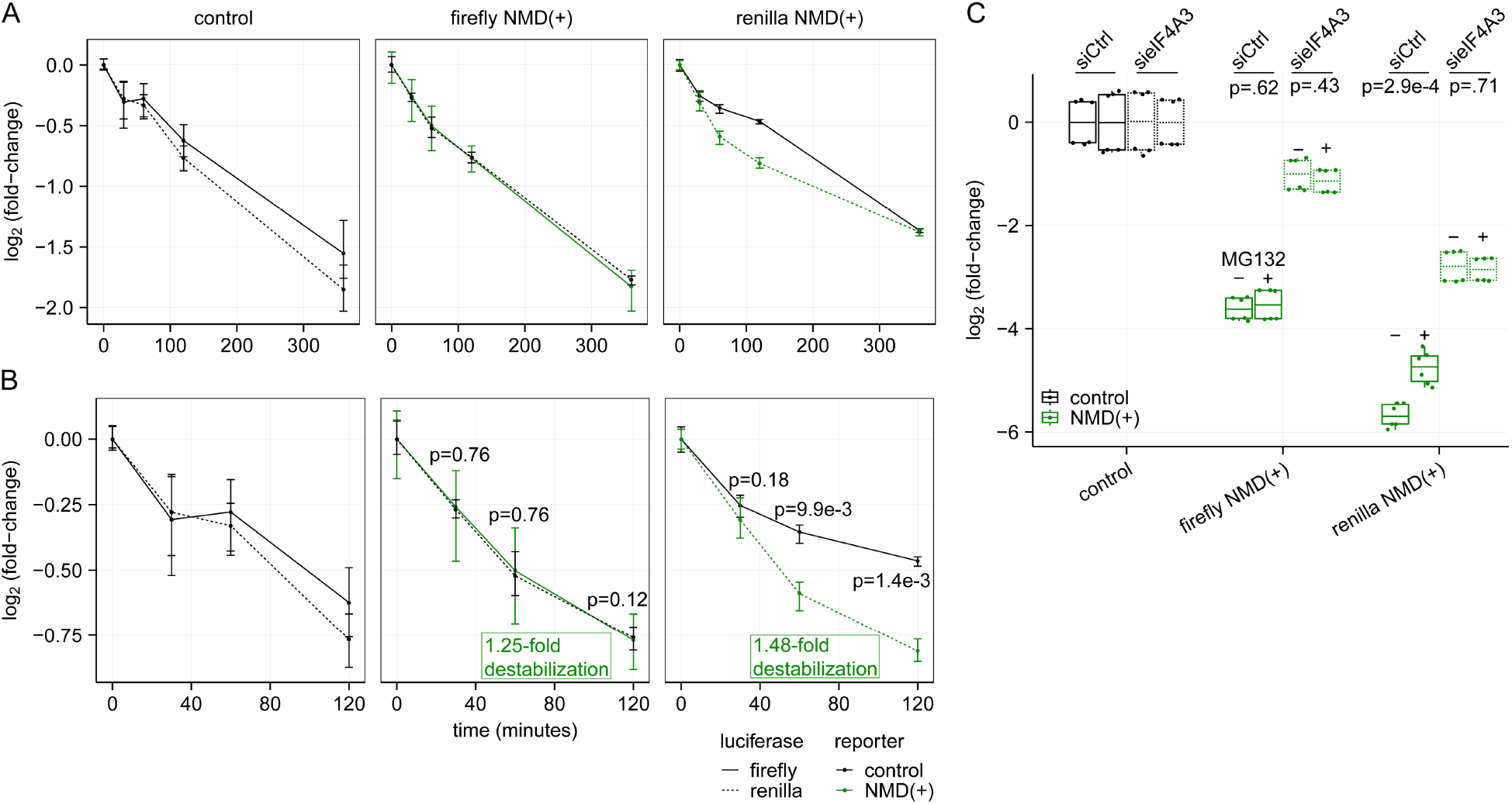
NMD(+) reporter proteins are degraded modestly faster than are control reporter proteins. (**A**) Line plots showing the decay kinetics of NMD reporter proteins after translation inhibition with cycloheximide. Each time point corresponds to n=3 technical replicates (n=2 biological replicates for the control panel, n=6 data points), with error bars showing the range of those values and the line plot connecting at the mean of the values. (**B**) Same as in (**A**), but only for the early time points. P-values were calculated as described for Figure 2B, using the ratio of firefly:renilla at each time point for the NMD(+) reporter cell lines compared to the control cell lines. The fold increase in destabilization/degradation of the NMD(+) reporter proteins relative to the corresponding control reporter proteins are based on the estimated half-lives of the reporter proteins calculated in Figure 4**—figure supplement 1B**. (**C**) Box plots showing NMD(+) reporter protein levels with and without MG132 treatment and eIF4A3 depletion relative to control reporter protein levels. An unpaired two-samples t-test was used for calculating the p-values.

Interestingly, the early time points do show faster degradation of NMD(+) reporter proteins relative to control reporter proteins (***Figure 4B***). However, the changes are relatively modest. To estimate the differences in half-lives of the reporter proteins at these early time points, we estimated best-fit exponential decay models (***Figure 4—figure supplement 1A***) and normalized to the control cell lines. The NMD(+) reporter proteins were degraded ∼1.1-1.6-fold faster than were the control reporter proteins (***Figure 4—figure supplement 1B***). In contrast, steady-state protein levels were ∼4-8-fold lower than were steady-state mRNA levels (***Figure 3B***), suggesting that increased degradation of proteins encoded by NMD-sensitive transcripts is not the primary mechanism underlying the marked protein-level suppression that we observed.

We next sought to determine if the modest increase in NMD(+) reporter protein decay that we observed was dependent on the ubiquitin-proteasome system. We treated our cell lines with MG132 to inhibit the proteasome and measured reporter protein levels. We observed no or very modest increases in NMD(+) reporter protein levels with MG132 treatment relative to no MG132 (***Figure 4C***, solid green boxes), consistent with modest increases in degradation rate (***Figure 4B***). Although the effects of MG132 treatment on protein levels were more notable for the renilla NMD(+) reporter than the firefly NMD(+) reporter, the effects for both were dwarfed by the effects of eIF4A3 depletion on protein levels (***Figure 4C***, dashed green boxes). Together, these data demonstrate that increased protein degradation is not the dominant mechanism leading to lower observed steady-state levels.

## DISCUSSION

We have developed a robust NMD reporter system for making precise, quantitative mRNA and protein level measurements (***Figure 1***). This system builds on previous reporters and adds numerous features, including (1) luciferase domains for high dynamic range protein-level measurements, (2) internal control reporters for accurate normalization across samples, (3) dox-inducibility for mRNA stability measurements, and (4) stable integration at the AAVS1 safe harbor loci for predictable genomic integration and uniform expression. The highly controlled nature of these reporters permitted us to clearly demonstrate that protein levels of the NMD(+) reporters were reduced to a greater degree than were mRNA levels, and to quantify the relative magnitude of mRNA- and protein-level suppression (***Figure 3***). Together with previous studies reporting protein-level suppression in both yeast and human cells, these findings imply that cells utilize mechanisms beyond mRNA decay to reduce the levels of potentially deleterious truncated proteins encoded by NMD-sensitive transcripts.

The modest increase in decay of the NMD(+) reporter proteins at early time points (***Figure 4B***) suggests that cells may have a mechanism to target truncated proteins for degradation, similar to the ribosome-associated protein quality control (RQC) pathway (*Joazeiro, 2019*). However, similar control and NMD(+) reporter protein levels at the late time point (***Figure 4A***) appear inconsistent with the existence of such a mechanism. One way to reconcile these differences is to hypothesize the presence of two populations of NMD transcripts: one consisting of transcripts that are rapidly degraded, and the other with transcripts that “escape” NMD and are degraded at a similar rate as control transcripts; only proteins derived from rapidly degraded transcripts are rapidly degraded themselves. This hypothesis is consistent with previous studies demonstrating the existence of two such pools of NMD-sensitive transcripts (*Cheng and Maquat, 1993*; Belgrader et al., 1994; Trcek et al., 2013; Kim et al., 2017; Hoek et al., 2019).

However, such a mechanism still would not fully explain the large difference between NMD(+) reporter mRNA and protein steady-state levels (***Figure 3B***), suggesting that additional mechanisms, such as reduced translation, likely modulate protein levels. Reduced translation of NMD-sensitive mRNAs has been observed in previous studies (Ishigaki et al., 2001; Chiu et al., 2004; *Sheth and Parker, 2006*; You et al., 2007; Isken et al., 2008; Lee et al., 2010; Kim et al., 2017) and, given the minimal protein decay differences that we observed (***Figure 4B***), is likely an important mechanism for limiting levels of proteins encoded by NMD-sensitive transcripts.

Our data suggest a model in which there are multiple layers of the NMD pathway, each of which acts to limit truncated protein accumulation (***Figure 5***). The first is the canonical, well-characterized mRNA degradation pathway, preventing truncated protein production by reducing the NMD-sensitive mRNA available to make proteins. The second is through limiting the accumulation of truncated proteins from the remaining mRNAs (via modestly increased protein degradation and reduced translation), leading to protein levels at a fraction of those of full-length proteins translated from NMD-insensitive transcripts. Neither mechanism is 100% efficient (some mRNAs escape NMD and some proteins are still translated from the remaining mRNAs), but the combination of both leads to an additive decrease in protein accumulation that prevents deleterious effects on cell health. A potential additional layer of modulation for truncated peptides encoded by endogenous NMD-sensitive mRNAs is inherent instability, which could lead to substantially faster degradation and even lower protein levels.

**Figure 5.**
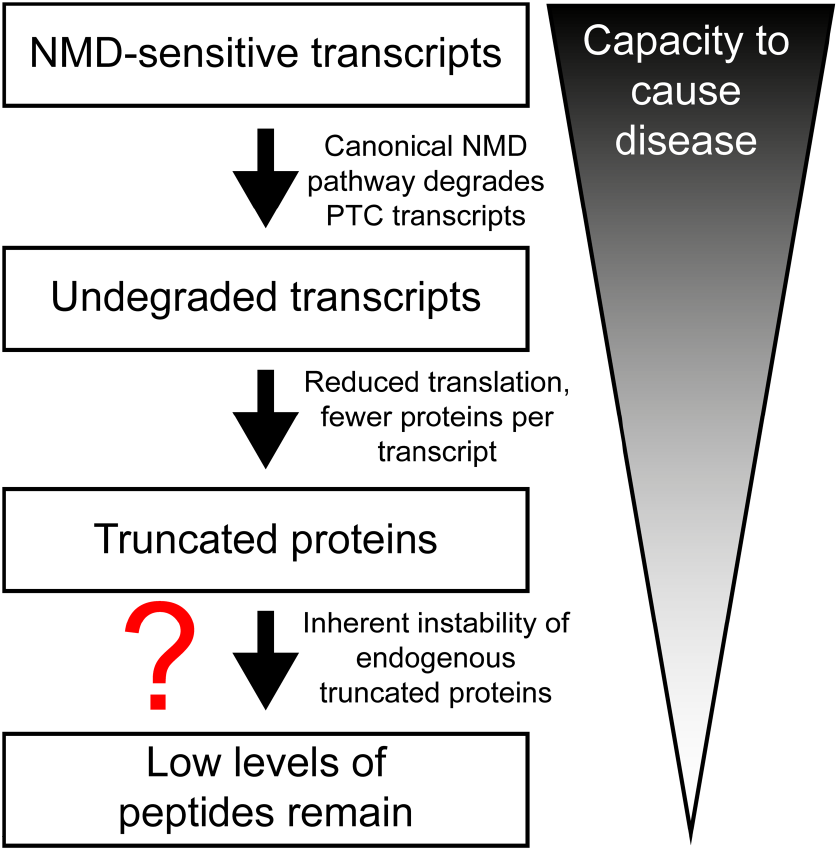
Model illustrating multiple layers of NMD pathway that complement one another to limit truncated protein accumulation.

The presumed purpose of NMD is to limit the accumulation of truncated proteins that could negatively affect cell health and homeostasis. The canonical mechanism for this is through recognition and degradation of NMD-sensitive mRNA, which is the well-characterized NMD pathway. Our data provide evidence for mechanisms complementary to the canonical pathway that further act to prevent the accumulation of truncated proteins. Future work is needed to determine if the limited protein accumulation from these NMD reporters is representative of most or all NMD-sensitive transcripts and their encoded proteins, and how other features – of both the transcript and peptide sequence – can influence this phenomenon.

## MATERIALS AND METHODS

### Design and cloning of luciferase-based NMD reporters

A set of firefly luciferase NMD reporters were obtained as a gift from Dr. James Inglese (Addgene IDs: 112085 and 112084). These reporters had the firefly luciferase sequence followed by either full-length or PTC39 beta-globin sequence in the p3xFLAG-CMV-10 backbone (Baird et al., 2018). To create renilla luciferase versions of these reporters, the plasmids were digested with EcoRI to cut on either side of the firefly luciferase sequence. The renilla luciferase sequence was amplified via PCR using two sets of primers with overhang sequences to facilitate isothermal assembly into the cut backbone (primer sequences listed in the Key Resources Table, RKB3257-3260). The renilla luciferase PCR amplicon (insert) and cut backbone were run on a 1% agarose gel and the DNA was isolated by gel extraction. The insert was then ligated into the cut backbone by isothermal assembly (NEBuilder® HiFi DNA Assembly, E2621L) according to the manufacturer’s protocol. The assembled plasmids were transformed into NEB^®^ Stable Competent *E. coli* (C3040H) and individual colonies were sequence verified.

The transient expression constructs described above were sub-cloned into donor plasmid backbones for CRISPR/Cas9 mediated genomic integration via homology directed repair. A backbone with homology arms to the AAVS1 safe harbor locus and with a dox-inducible promoter (Natsume et al., 2016; Addgene ID 72835) was cut with MluI and BglII. The luciferase-beta-globin sequence was amplified via PCR using primers with overhang sequences (RKB3454-3455) to facilitate isothermal assembly into the cut backbone. The cloning proceeded as described above, with the final assembled plasmids sequence verified. All of these plasmids are available on Addgene (see Key Resources Table).

### Cell culture and genome engineering of HEK 293 cells

Flp-In™ T-REx™ 293 cells (ThermoFisher, R78007) were cultured at 37°C and 5% CO_2_ in DMEM media with 10% FBS and 1% penicillin-streptomycin and split every 2-3 days before reaching full confluence. For stable integration of the NMD reporters, 293 cells were plated at 40% confluency in a well of a 12-well plate the day before transfecting. For each well, ∼400 ng of firefly donor plasmid, ∼400 ng of renilla donor plasmid, and ∼400 ng of Cas9/AAVS1-sgRNA expressing plasmid were used. The DNA and 2.4 µL of P3000 reagent were diluted in 60 µL of Opti-MEM (Gibco) and separately 1.8 µL of Lipofectamine 3000 Reagent (Invitrogen, L3000-015) was diluted in 60 µL Opti-MEM. The transfections proceeded according to the manufacturer’s protocol, with 100 µL of transfection mix added to each well already containing 1 mL of media.

One day after transfecting, puromycin (Gibco, A11138-03) was added (2 µg/mL) to select for successfully transfected cells. The next day, cells were split from each well into a 10 cm plate and grown in puromycin-containing media for several days to select for stable integration of the reporters. Polyclonal cell lines were cryopreserved and used for subsequent experiments.

### Induction of reporters and depletion of NMD factors

Cells were plated at 20% confluency in poly-L-lysine coated wells of a 12-well culture plate the day before siRNA transfections. For each well, 3 µL of Lipofectamine RNAiMAX Reagent (Invitrogen, 13778-150) was diluted in 50 µL of Opti-MEM (Gibco, 31985-062) and separately siRNA was diluted in 50 µL of Opti-MEM. The transfections proceeded with the manufacturer’s protocol, with 100 µL of transfection mix added to each well already containing 900 µL of media. The final concentration of siRNA for each well was 20 nM. The cells were collected 72 hours after transfection. To induce expression of the stably integrated reporters, doxycycline (Sigma, D9891) was added to a final concentration of 1 µg/mL to each well 24 hours before cell collection.

### RNA extraction and RT-PCR to confirm expected splicing of reporter mRNA

Cell pellets were lysed with TRIzol Reagent (Invitrogen, 15596-026) and processed according to the manufacturer’s protocol for RNA isolation. The RNA pellet was resuspended in 50 µL of water and then subjected to RNeasy (Qiagen, 74104) column purification and DNase digestion. The 50 µL of RNA was added to 300 µL RLT lysis buffer, mixed with 350 µL 70% ethanol, and transferred to an RNeasy spin column. The samples were then processed according to the manufacturer’s protocol using RNase-Free DNase Set (Qiagen, 79254) for on-column DNase digestion.

The purified RNA was confirmed to be free of DNA contamination using end-point PCR with primers specific to the firefly luciferase sequence (RKB2250-2251). cDNA was synthesized from 3 µg of RNA per sample using SuperScript IV Reverse Transcriptase (Invitrogen, 18090050) using oligo-dT primers according to the manufacturer’s protocol. To check splicing, forward primers were designed in either the firefly or renilla sequence and reverse primers were designed in the 3’-UTR sequence just downstream of the last beta-globin exon. End-point RT-PCR was performed on the cDNA samples using these primers (RKB3600-3612, **Figure 1—figure supplement 2**), and the PCRs were run on 1% agarose gels to visualize the size of the PCR amplicons.

### NMD reporter mRNA steady-state level measurement

The cDNA reactions were diluted 1:50 and 4 µL was used per 10 µL qRT-PCR reaction with PowerUp SYBR Green Master Mix (ThermoFisher, A25742) and final primer concentrations of 500 nM in 384-well plates (ThermoFisher, AB1384). Two unique primer sets (Key Resources Table) were used for each luciferase and two reference genes were also quantified with qRT-PCR using an ABI QuantStudio 5 Real-Time PCR System (ThermoFisher). Three technical replicate reactions were performed for each unique primer set for each sample.

To quantify the reporter mRNA levels from the qRT-PCR data, the means of the technical replicates for each reporter primer set were used. Each of the two firefly primer sets was compared to each of the two renilla primer sets, for a total of n=4 normalized technical replicates in each sample. Those replicates were normalized to replicates from two control cell lines for a total of eight data points, all plotted relative to the control cell lines.

### Decay kinetics of NMD reporter mRNA

Cells were plated at 10% confluency in poly-L-lysine coated wells of a 12-well culture plate the day before induction. The following day the media was replaced with doxycycline-containing media 24 hours before cell collection. To turn off reporter expression to measure mRNA decay kinetics, doxycycline containing media was removed and the cells were washed with PBS and standard media (no dox) before being replaced with standard media for a specified length of time before the cells were harvested. RNA extraction, cDNA synthesis, and qRT-PCR were performed as described above.

For examining reporter mRNA decay kinetics from qRT-PCR data, each of the luciferase primer sets (two sets for firefly and two sets for renilla) was normalized to two reference genes (*RPL27* and *SRP14*) for a total of n=4 technical replicates for each luciferase in each sample. Those values were normalized to the sample collected at time 0 for each cell line and plotted relative to time 0 to show the decrease in reporter mRNA over time.

### Western blotting for NMD reporter proteins

Cells were collected from individual wells of a 12-well culture plate, centrifuged at 1200 rpm for 5 min, washed with PBS, and lysed using 50 µL of NP40 Cell Lysis Buffer (Invitrogen, FNN0021) supplemented with protease and phosphatase inhibitors (Pierce, A32955). Lysates were sonicated, incubated on ice, and spun down at 10,000 x g at 4°C for 15 min; the supernatant was collected for downstream assays. The protein concentration was quantified using the Qubit Protein Assay (Thermo, Q33212). 12 µg of protein per sample was used for gel electrophoresis with a NuPAGE™ 4-12% Mini Protein Gel (Invitrogen, NP0323) in a Mini Gel Tank (Invitrogen) with 1X NuPAGE™ MOPS SDS Running Buffer (Invitrogen, NP0001). After electrophoresis, protein was transferred overnight to a nitrocellulose membrane (Invitrogen, LC2001) using 1X NuPAGE™ Transfer Buffer (Invitrogen, NP0006-1) with 10% methanol. After transfer, protein bands were visualized with Ponceau stain (Sigma, P7170) to confirm protein transfer and even loading.

The membrane was blocked with Odyssey Blocking Buffer (PBS) (LI-COR Biosciences, 927-40000) for one hour at 4°C with gentle shaking. The blot was probed with primary antibodies overnight at 4°C with gentle shaking; specific antibodies and dilutions used are listed in the Key Resources Table. Following overnight incubation, the blot was washed 3 times for 5 min with 1X TBST buffer at room temperature with gentle shaking. The blot was then probed with IRDye® secondary antibodies (LI-COR Biosciences) for 1 hour at room temperature with gentle shaking, followed by a final sequence of washes as described above. The blot was imaged on an Odyssey® CLx Imaging System and images were processed using Fiji (ImageJ v2.1.0).

The western blot did not show full depletion of the eIF4A3 protein. However, the immunogen sequence used to generate the eIF4A3 antibody is highly conserved among eIF4A3, eIF4A1, and eIF4A2, and all three proteins are similar size. We predict that the antibody is likely also binding eIF4A1 and eIF4A2 on the western blot and the band shown corresponds to all three of those proteins. Given that the bands have modestly reduced intensity and the NMD reporter protein levels increase in the lanes with sieIF4A3 samples, we concluded that the siRNAs are likely effectively depleting eIF4A3.

### Dual-luciferase assay for NMD reporter protein level measurement

The Dual-Luciferase® Reporter Assay System (Promega, E1910) was used for measuring luciferase levels from NMD reporter cell lines according to the manufacturer’s protocol. Briefly, individual wells of a 12-well culture plate were washed with 1 mL of PBS before 200 µL of 1X Passive Lysis Buffer was added directly to each well. The culture plates were placed on an orbital shaker with gentle rocking for 15 minutes at room temperature to achieve complete lysis. Cell lysates were collected and centrifuged at max speed for 30 seconds and the supernatants were collected and used for subsequent assays.

The dual-luciferase assay was performed using a Cytation™ 5 plate reader with a dual-injector system (BioTek). For each sample, 20 µL of lysate was transferred to wells of a 96-well plate. The plate reader was set-up and programmed to inject 100 µL of Luciferase Assay Reagent II (LAR II) from the first injector and 100 µL of Stop & Glo® Reagent from the second injector. Timing for measuring luminescence was set according to the Dual-Luciferase® Reporter Assay System protocol.

### Monoclonal NMD reporter cell line generation

Polyclonal cell lines underwent single cell sorting into 96-well culture plates using an MA900 multi-application cell sorter (Sony Biotechnology). Single cells were grown in 50 µL of DMEM media supplemented with 20% FBS per well, with an additional 50 µL of media added every 3-4 days to maintain optimal growth conditions for cells at low confluence. Cells began reaching confluence in individual wells 2-3 weeks after sorting, at which time the cells were split into 24-well plates. Upon reaching confluence in the 24-well plates, cells were split into two separate 12-well plates: one to continue propagating cells and another for a dual-luciferase assay to determine which luciferase reporters were stably integrated. Monoclonal lines with luciferase expression were cryo-preserved and a subset of cells from each line were collected for gDNA extraction with DNeasy® Blood & Tissue Kit (Qiagen, 69504) to confirm stable integration of the reporters at the AAVS1 loci via gDNA-PCR.

A forward primer was designed in the AAVS1 sequence outside of the left homology arm on the donor plasmid (RKB2392), while the reverse primer was designed in the luciferase sequence (**Figure 3—figure supplement 1**) such that only stable integration of the reporters at that locus would yield a PCR amplicon. Separate reverse primers were designed for firefly and renilla luciferase sequences (RKB3517 and RKB3531), and monoclonal lines with amplicons specific to both luciferases were used for subsequent experiments.

### Proteasome inhibition in NMD reporter cell lines

Reporter cell lines were grown in 12-well culture dishes and treated with siRNAs as described above. Cells were treated with a final concentration of 10 µM MG132 (Sigma, C2211) to inhibit the proteasome; control samples were treated with DMSO (vehicle) (Sigma, D2650). Cells were lysed in 200 µL of 1X Passive Lysis Buffer (dual-luciferase assay) 6 hours after MG132 addition. The samples were processed for use in the dual-luciferase assay as described above.

### Decay kinetics of NMD reporter proteins

Cells were grown in 12-well culture plates and reporter expression was induced with doxycycline as described above. To inhibit translation, cells were treated with cycloheximide (Sigma, C7698) at a final concentration of 100 µg/mL for specified lengths of time before being lysed with 200 µL of 1X Passive Lysis Buffer. The samples were processed for use in the dual-luciferase assay.

For examining reporter protein decay kinetics from the dual-luciferase assay data, the luminescence values for each luciferase were plotted relative to the time 0 value to show the change in reporter protein levels over time after translation inhibition. The half-lives of the reporters were calculated using linear regression of the mean of the technical replicates at each time point for each reporter. The half-lives of the NMD(+) reporter proteins were normalized to those of the control reporter proteins and control reporter cell line to get the “fold destabilization” (**Figure 4B**) relative to the control reporter.

## ACKNOWLEDGEMENTS

We thank members of the Bradley lab for helpful discussions; former lab members Qing Feng, Sujatha Jagannathan, and Heather Johns for technical help with NMD reporters and guidance with project directions; Robert Hogg, James Inglese, and Ken Cheng for sharing plasmids (Addgene IDs 112084 and 112085) and cells from Baird et al., 2018 (pKC-4.06); and Arvind Subramaniam for providing feedback on the manuscript. RKB was supported in part by the NIH/NHLBI (R01 HL151651); NIH/NCI (R01 CA251138); NIH/NHLBI (R01 HL128239); NIH/NIDDK (R01 DK103854); Blood Cancer Discoveries Grant program through the Leukemia & Lymphoma Society, Mark Foundation for Cancer Research, and Paul G. Allen Frontiers Group (8023-20); and Dept. of Defense Breast Cancer Research Program (W81XWH-20-1-0596). RKB is a Scholar of The Leukemia & Lymphoma Society (1344-18) and holds the McIlwain Family Endowed Chair in Data Science. This research was supported in part by the NIH/NCI (Cancer Center Support Grant P30 CA015704). DBU was supported in part by NIH/NIGMS T32 GM007270.

## AUTHOR CONTRIBUTIONS

DBU and RKB designed the study. DBU performed experiments and analyzed data. DBU and RKB wrote the manuscript.

## DATA AVAILABILITY

All source data is available as source data files.

---

**Figure 1—figure supplement 1.**
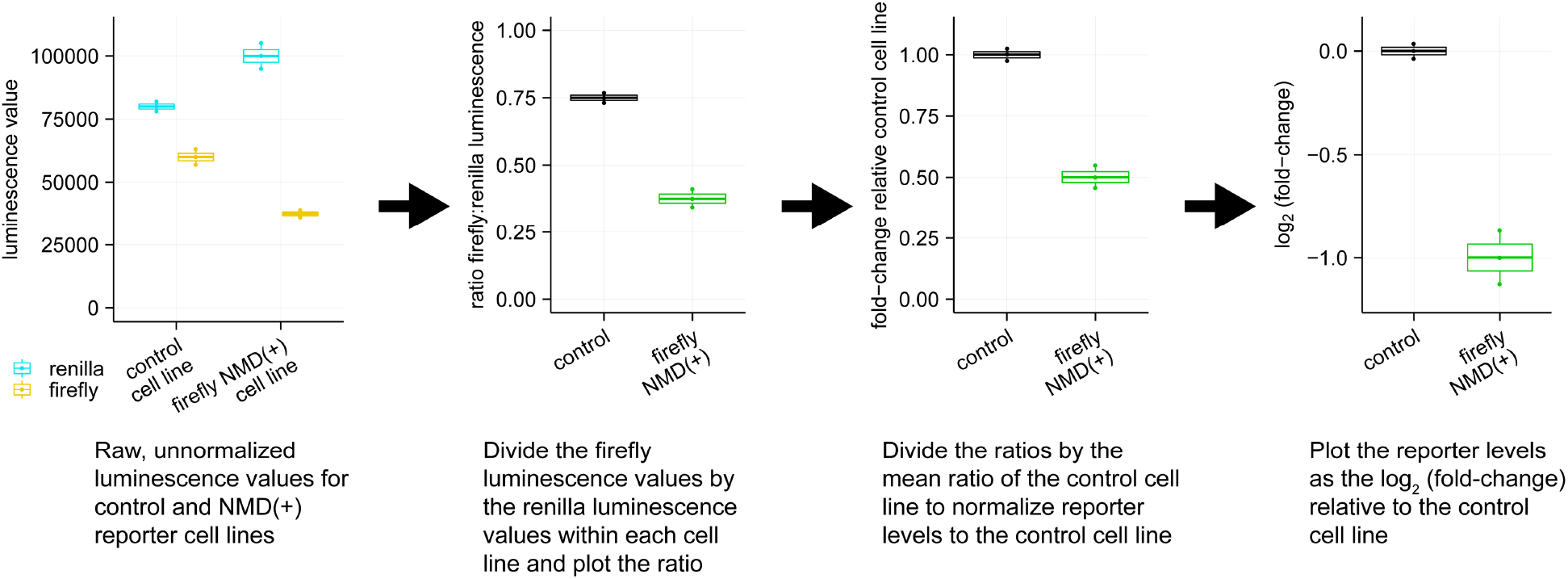
Mock dual-luciferase assay data demonstrating how to normalize the luminescence values of the luciferases and how to plot the level of the NMD(+) reporter relative to the control reporter.

**Figure 1—figure supplement 2.**
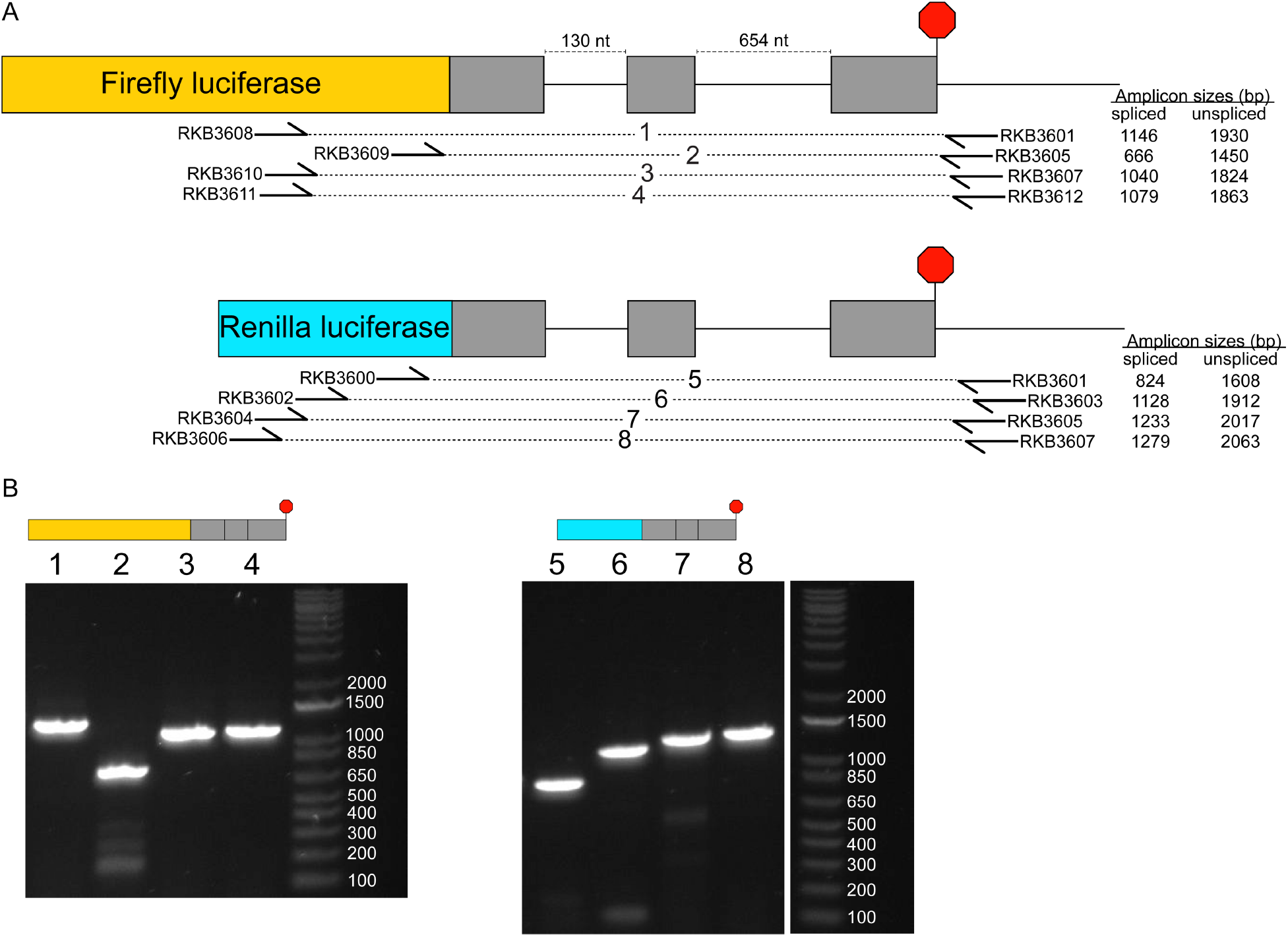
Stably integrated luciferase reporters are spliced correctly. (**A**) Schematics showing how primers were designed for each luciferase reporter to confirm correct splicing. The forward primers were designed in the luciferase sequence while the reverse primers were designed in the 3’-UTR; if either intron were not spliced out there would be a noticeable size shift in the PCR amplicon. The sizes are shown for the expected PCR amplicons for each primer set from either the fully spliced reporter mRNA or the fully unspliced reporter mRNA. The primer sequences are listed in the Key Resources Table. (**B**) End-point RT-PCR using the primer sets shown in (**A**) and cDNA from cell lines with stably integrated luciferase reporters. The number above each lane corresponds to the primer pair used for that PCR (listed in (**A**)).

**Figure 1—figure supplement 3.**
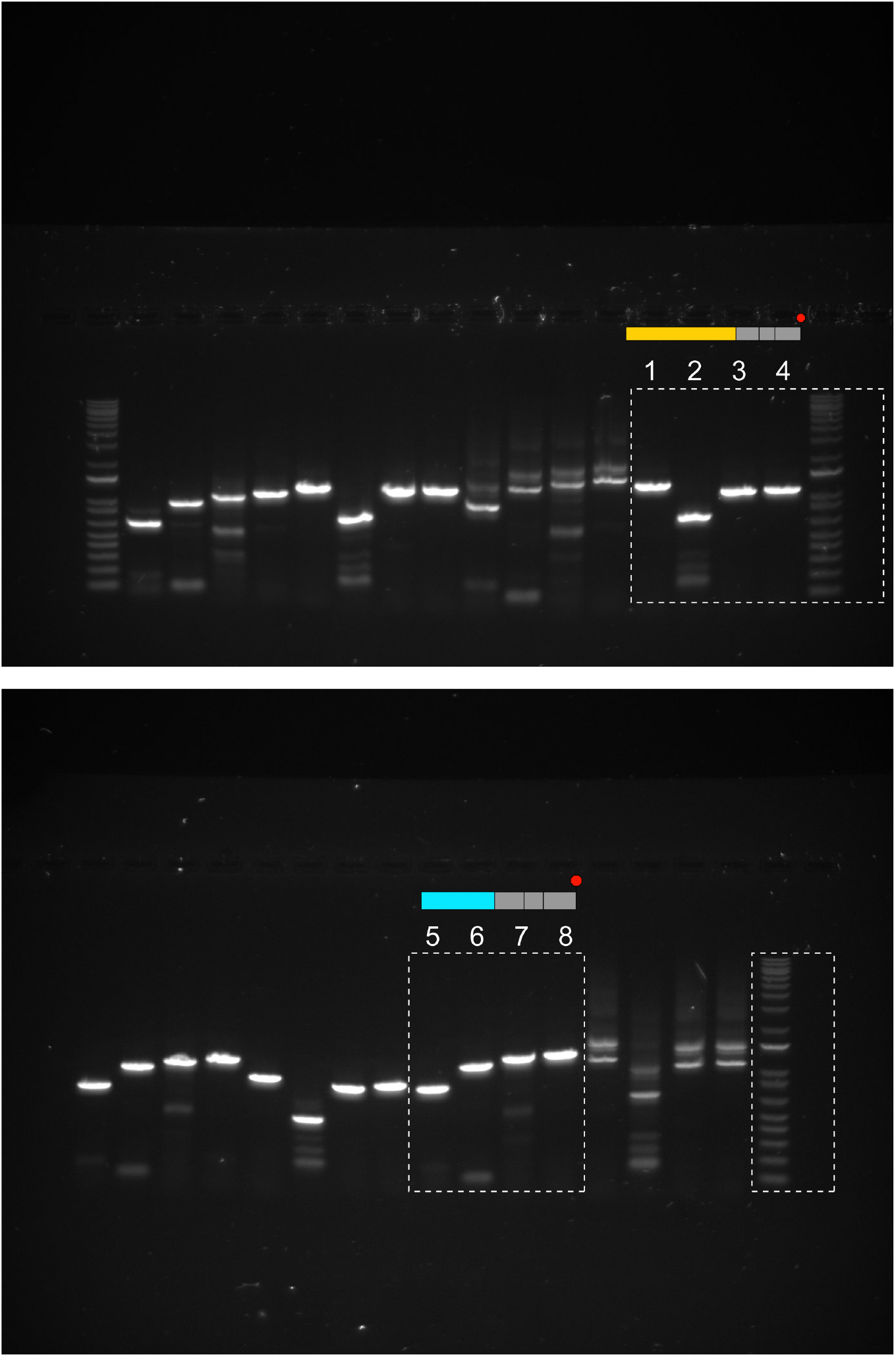
Raw images of the RT-PCR gels shown in Figure 1**—figure supplement 2**. The cropped regions are shown in the dashed boxes.

**Figure 2—figure supplement 1.**
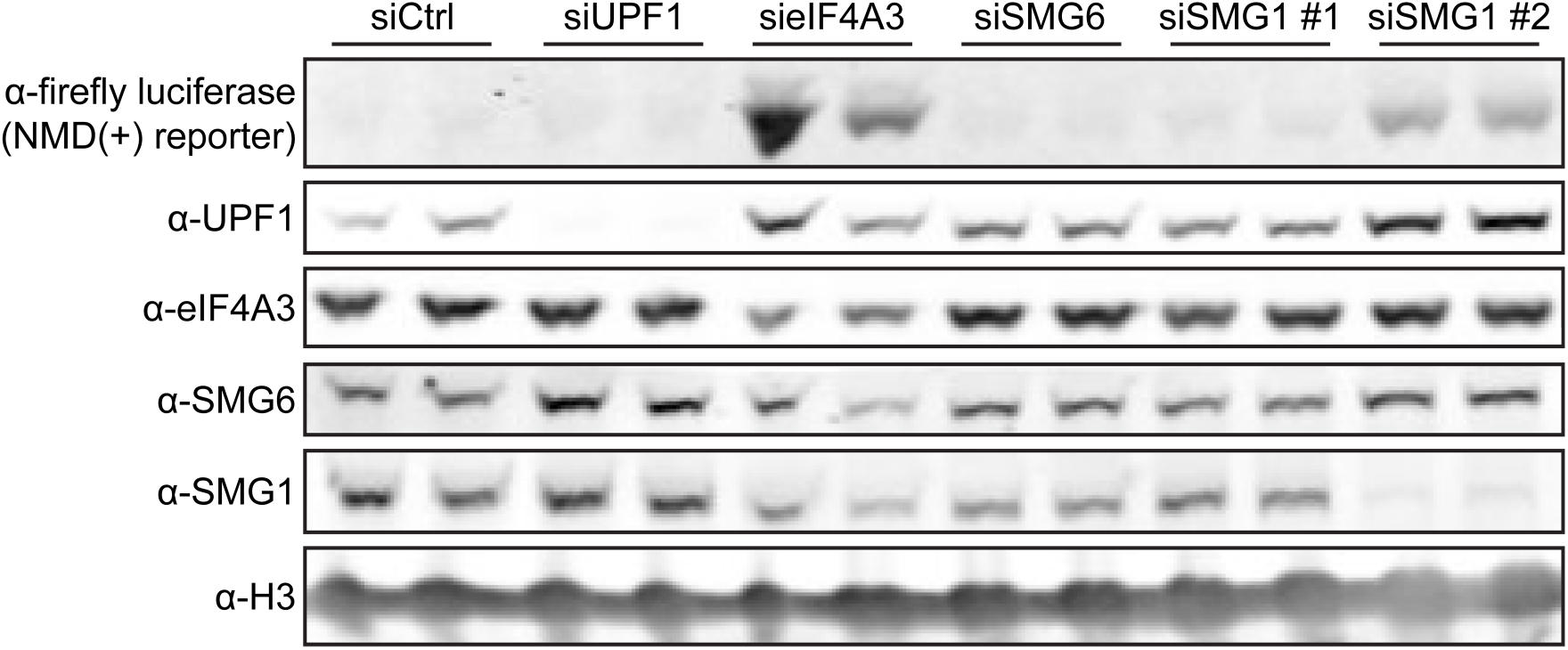
Western blot showing the depletion of several factors involved in NMD and the corresponding change in firefly NMD(+) reporter protein levels in samples from HEK-293 cells. For each pair of siRNA samples, the first lane corresponds to 20 nM siRNA concentration and the second lane corresponds to 50 nM siRNA concentration. Subsequent experiments used 20 nM siRNA concentrations.

**Figure 2—figure supplement 2.**
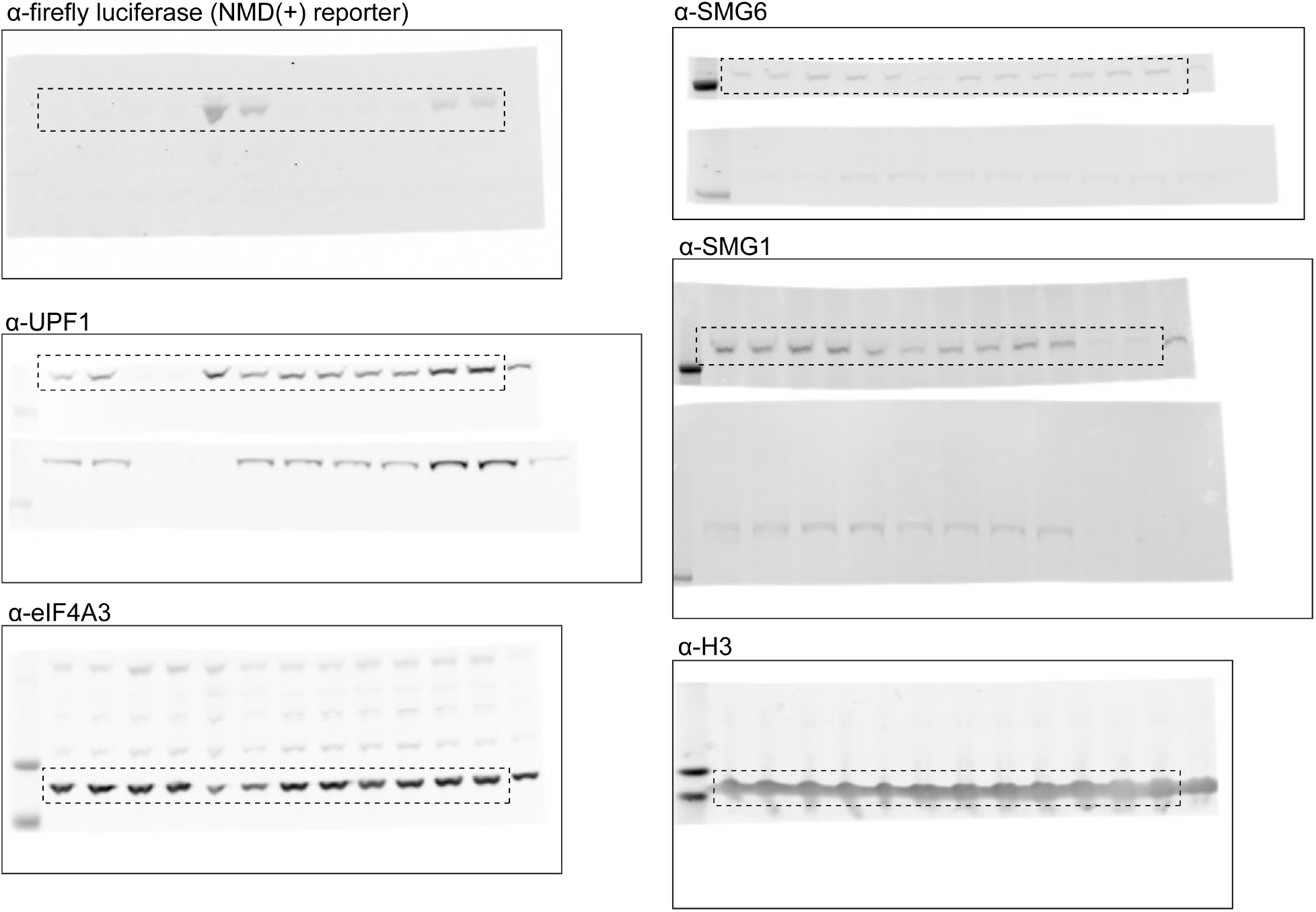
Raw images of the western blots shown in Figure 2**—figure supplement 1**. The cropped regions are shown in the dashed boxes.

**Figure 3—figure supplement 1.**
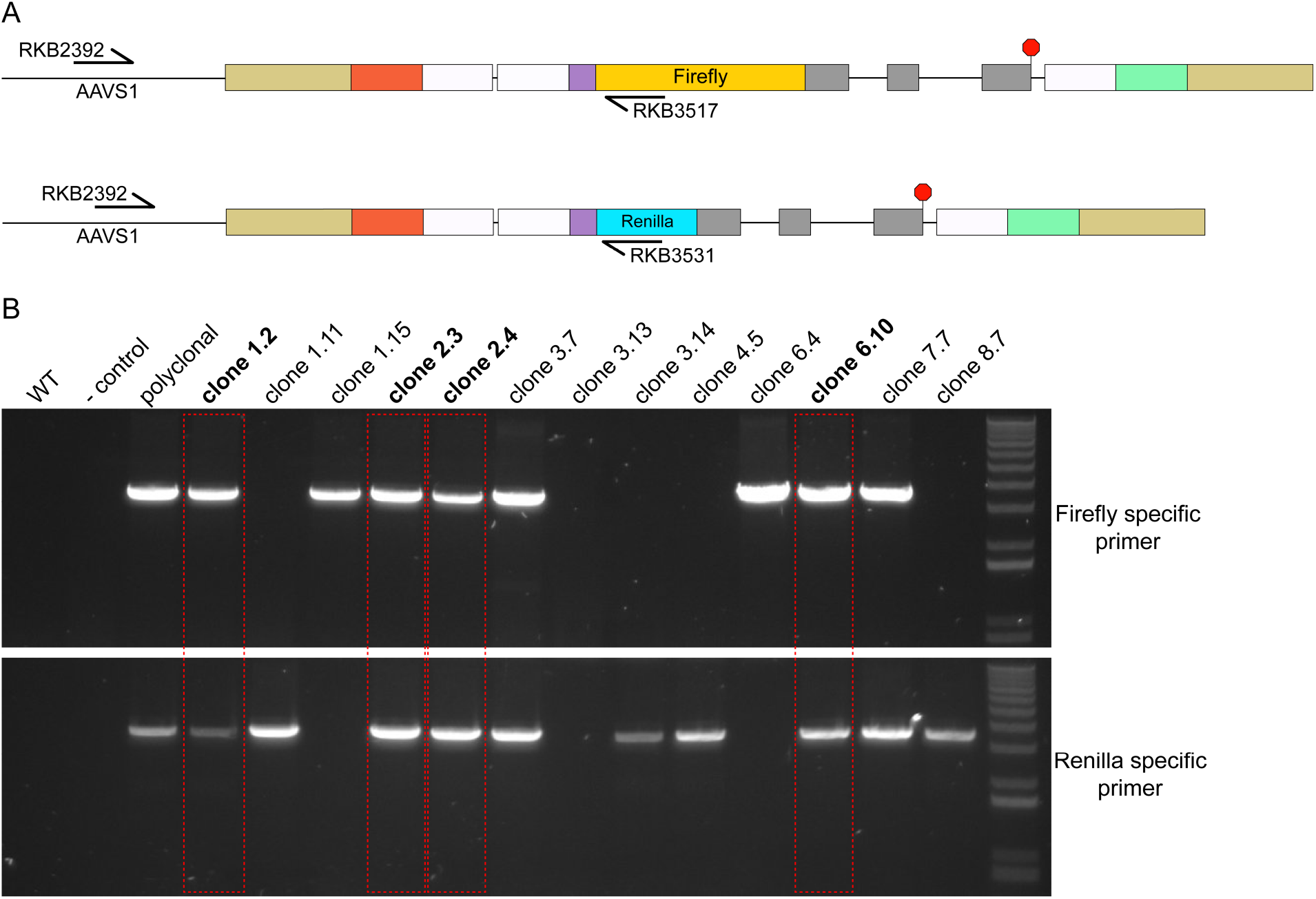
Monoclonal cell lines have both luciferase reporters integrated at the AAVS1 loci. (**A**) Schematic showing primer design to confirm integration of both reporters at the AAVS1 loci. The forward primer was designed upstream of the AAVS1 homology arm region, while the reverse primer was designed inside the luciferase sequence. The different primers for the different luciferase sequences allow us to distinguish whether the firefly and/or renilla reporter is stably integrated. Primer sequences are listed in the Key Resources Table. (**B**) Genomic DNA PCR using the primer sets shown in (**A**). The bands outlined in the red boxes correspond to the clonal cell lines that had stable integration of both luciferase reporters at the AAVS1 loci and were subsequently used for dual-luciferase assay measurements.

**Figure 3—figure supplement 2.**
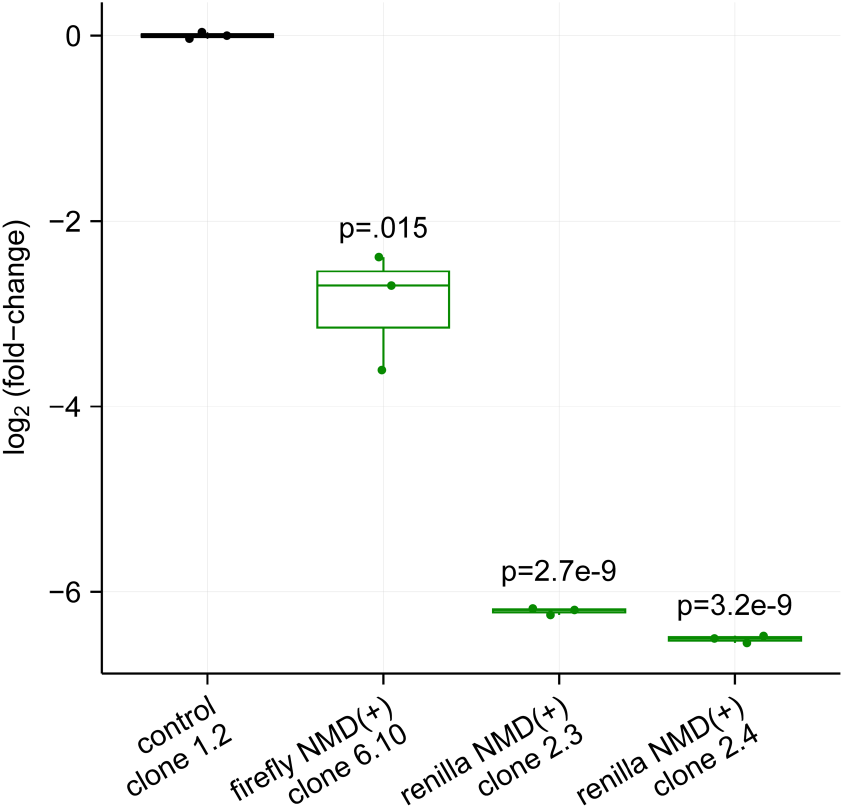
Box plots showing the NMD reporter protein levels from monoclonal cell lines with both firefly and renilla luciferase reporters stably integrated in the AAVS1 loci. The levels are normalized to the control reporter monoclonal cell line. Each box plot shows n=3 technical replicates normalized to n=1 biological replicate. An unpaired two-samples t-test was used for calculating the p-values, which correspond to the comparison between the monoclonal control cell line and each monoclonal NMD(+) reporter cell line.

**Figure 3—figure supplement 3.**
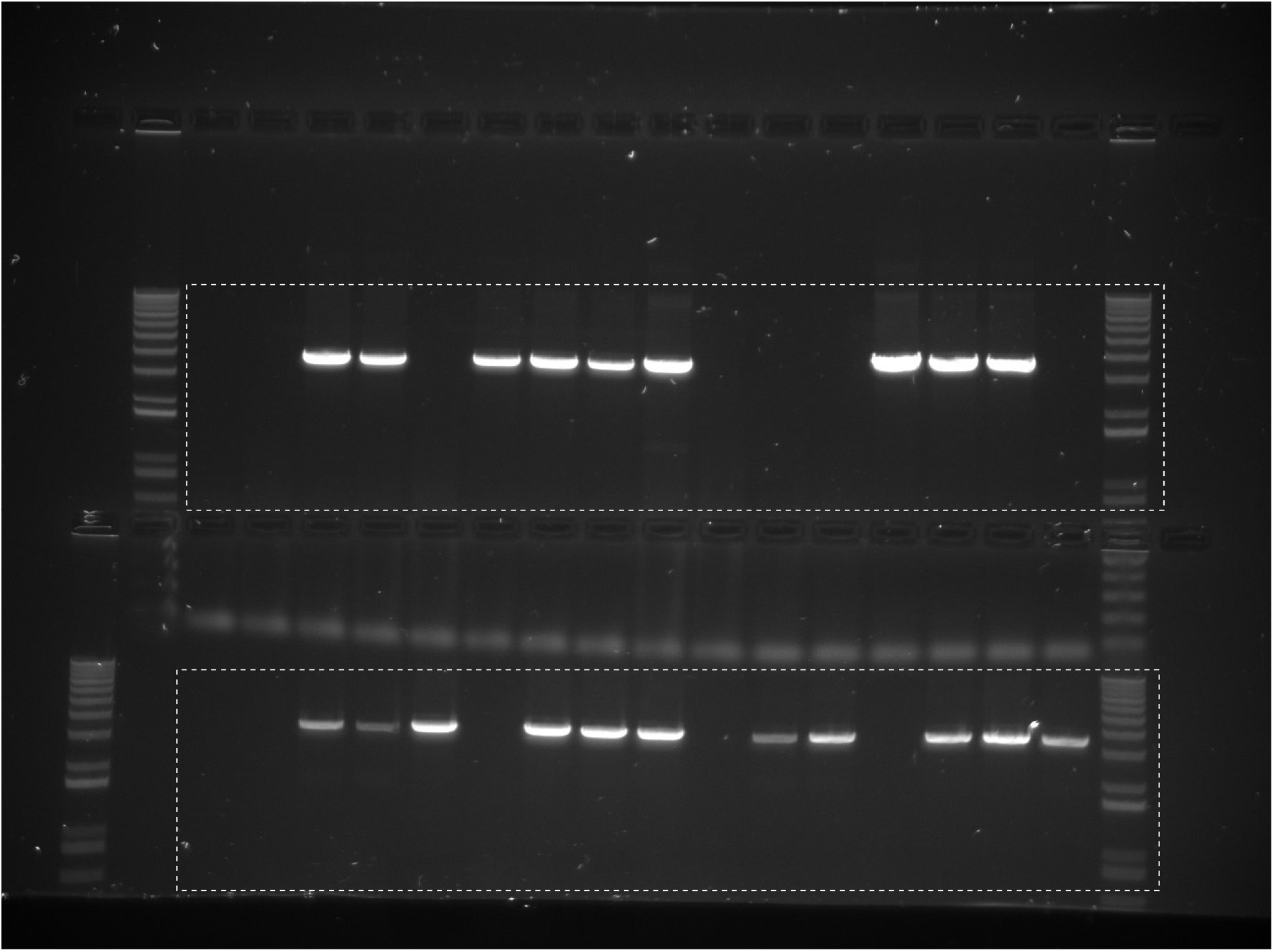
Raw image of the gDNA PCR gels in Figure 3**—figure supplement 1**. The cropped regions are shown in the dashed boxes.

**Figure 4—figure supplement 1.**
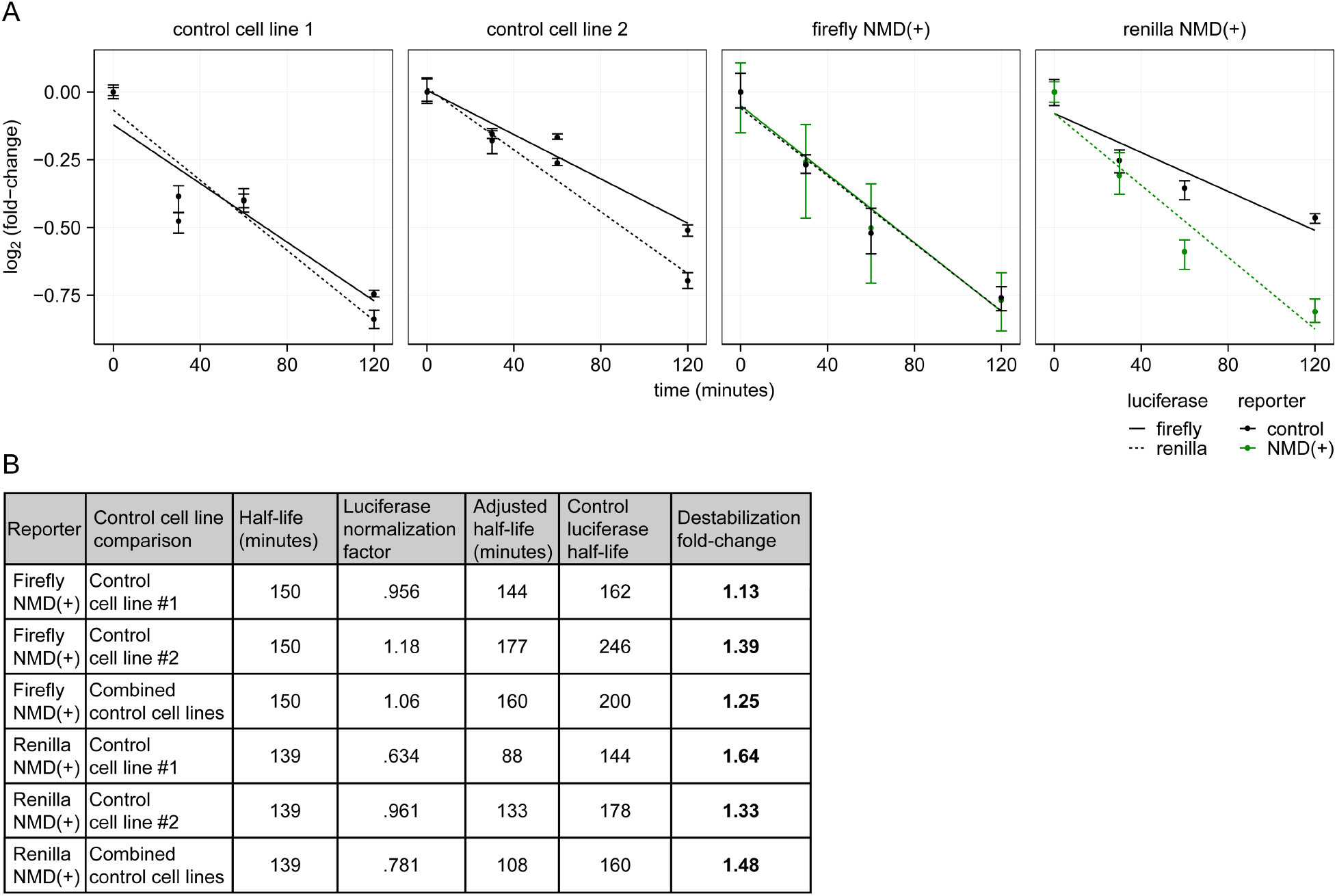
Estimating the half-lives of the NMD reporter proteins. (**A**) Same data as in Figure 4B, but with a best-fit line plotted as calculated from linear regression of the mean of the technical replicates at each time point for each reporter. The best fit line was used to calculate the unnormalized half-life for each reporter protein in each cell line based on the time (x-axis) at which the best-fit line crosses the 50% protein level (relative to time 0). The two control cell lines are plotted separately. (**B**) Table showing fold-change in destabilization of the NMD(+) reporter protein relative to the control reporter protein for each luciferase reporter. The half-lives of the reporter proteins in the NMD(+) cell lines were normalized to the half-lives of the reporter proteins in the control cell lines (using the half-lives of the control luciferase reporter proteins that both cell lines have in common) in order to more accurately estimate the differences in half-lives. The half-life of the control luciferase in the control reporter cell line was divided by the half-life of the corresponding control luciferase in the NMD(+) reporter cell line (e.g. control reporter cell line firefly/NMD(+) reporter cell line firefly) to get the “luciferase normalization factor.” The half-life of the NMD(+) luciferase in the NMD(+) reporter cell line was then multiplied by the luciferase normalization factor to get the “adjusted half-life.” The half-life of the corresponding control luciferase in the control reporter cell line was then divided by the adjusted half-life of the NMD(+) luciferase in the NMD(+) reporter cell line (e.g. control reporter cell line renilla half-life/NMD(+) reporter cell line renilla adjusted half-life) to get the fold-change in destabilization of the NMD(+) reporter protein relative to the control reporter protein. For each NMD(+) reporter luciferase, the comparison was made with each control cell line separately and the two control cell lines combined (“Control cell line comparison” column).

## APPENDIX

### Key Resources Table

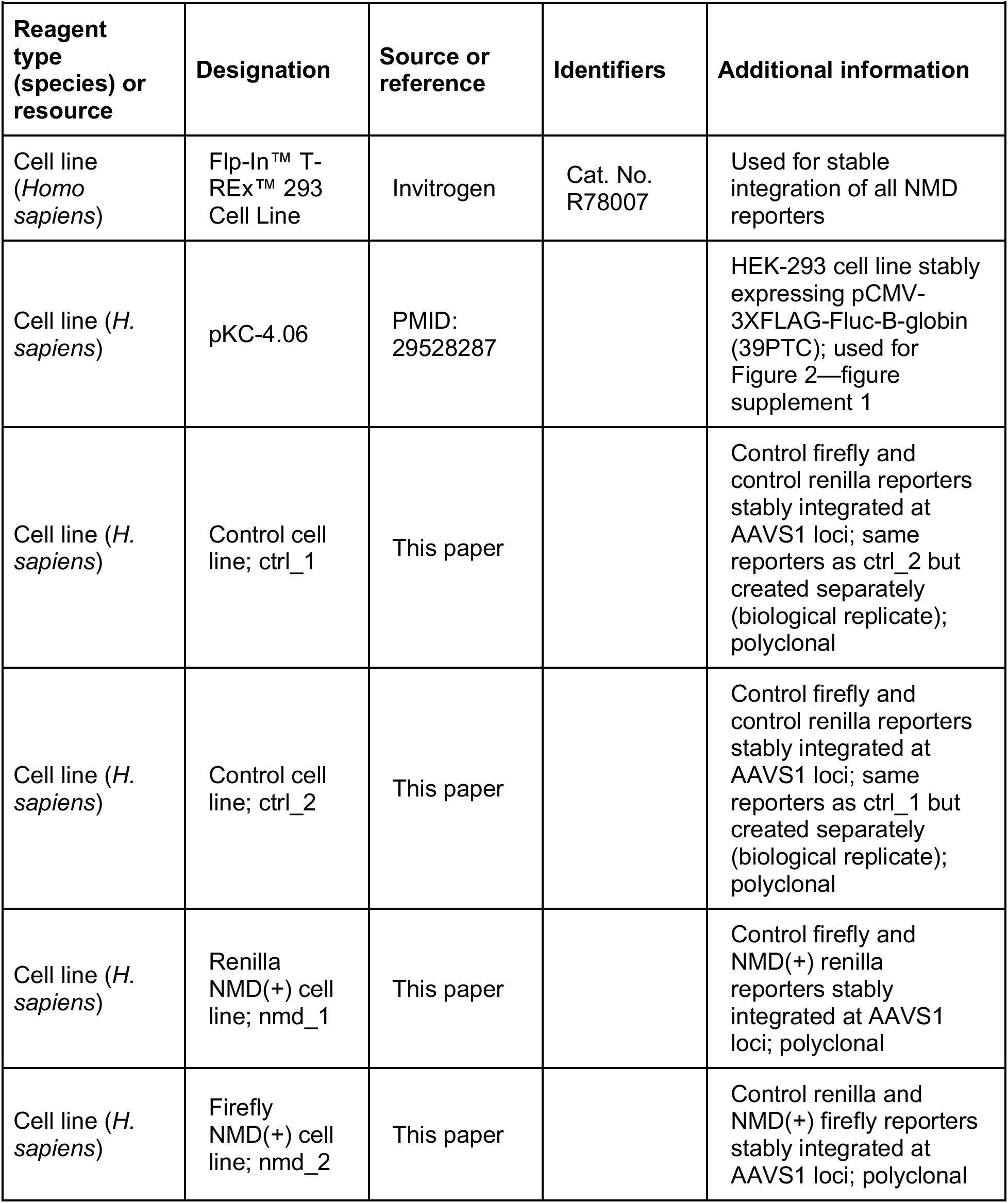
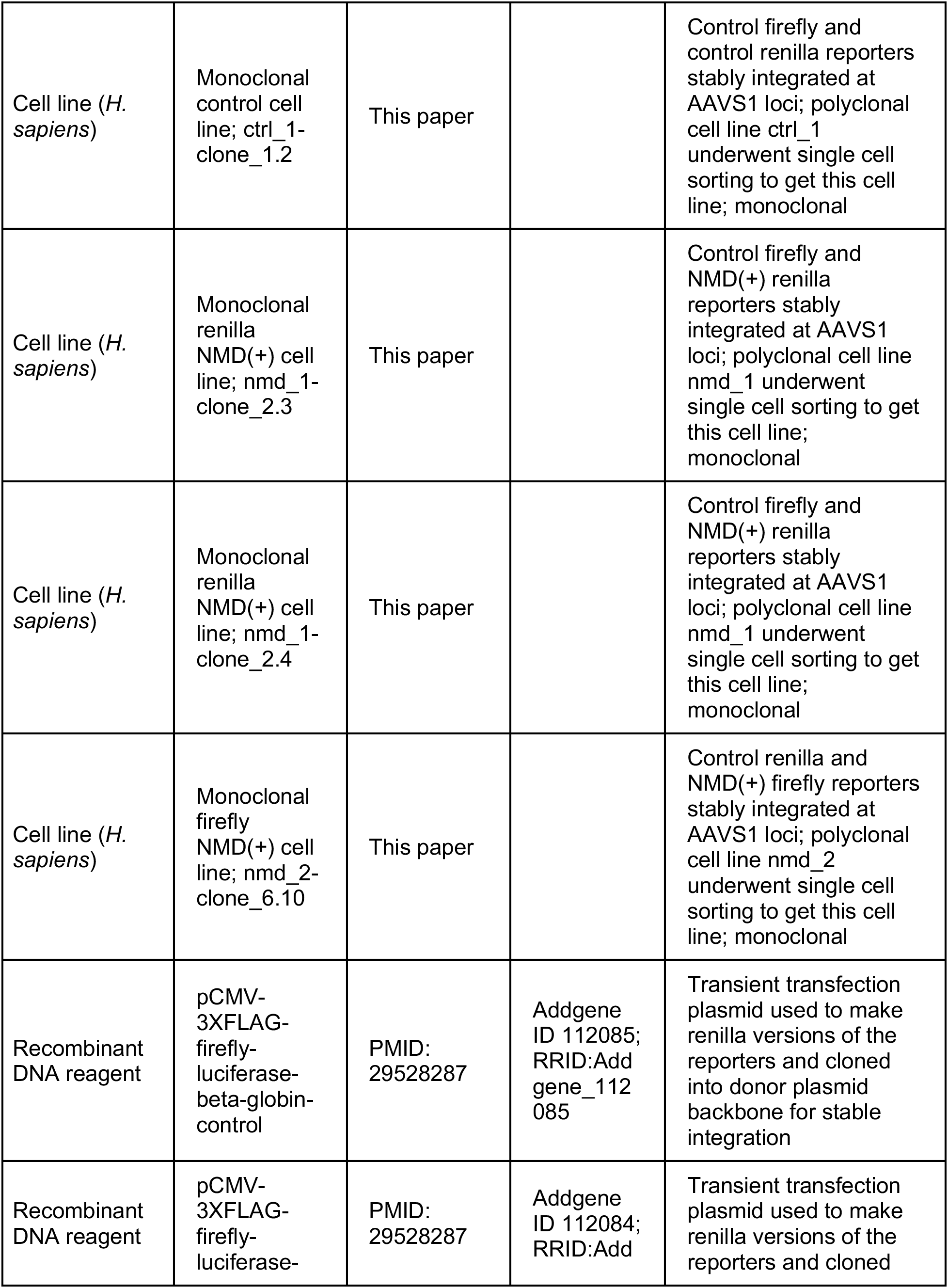
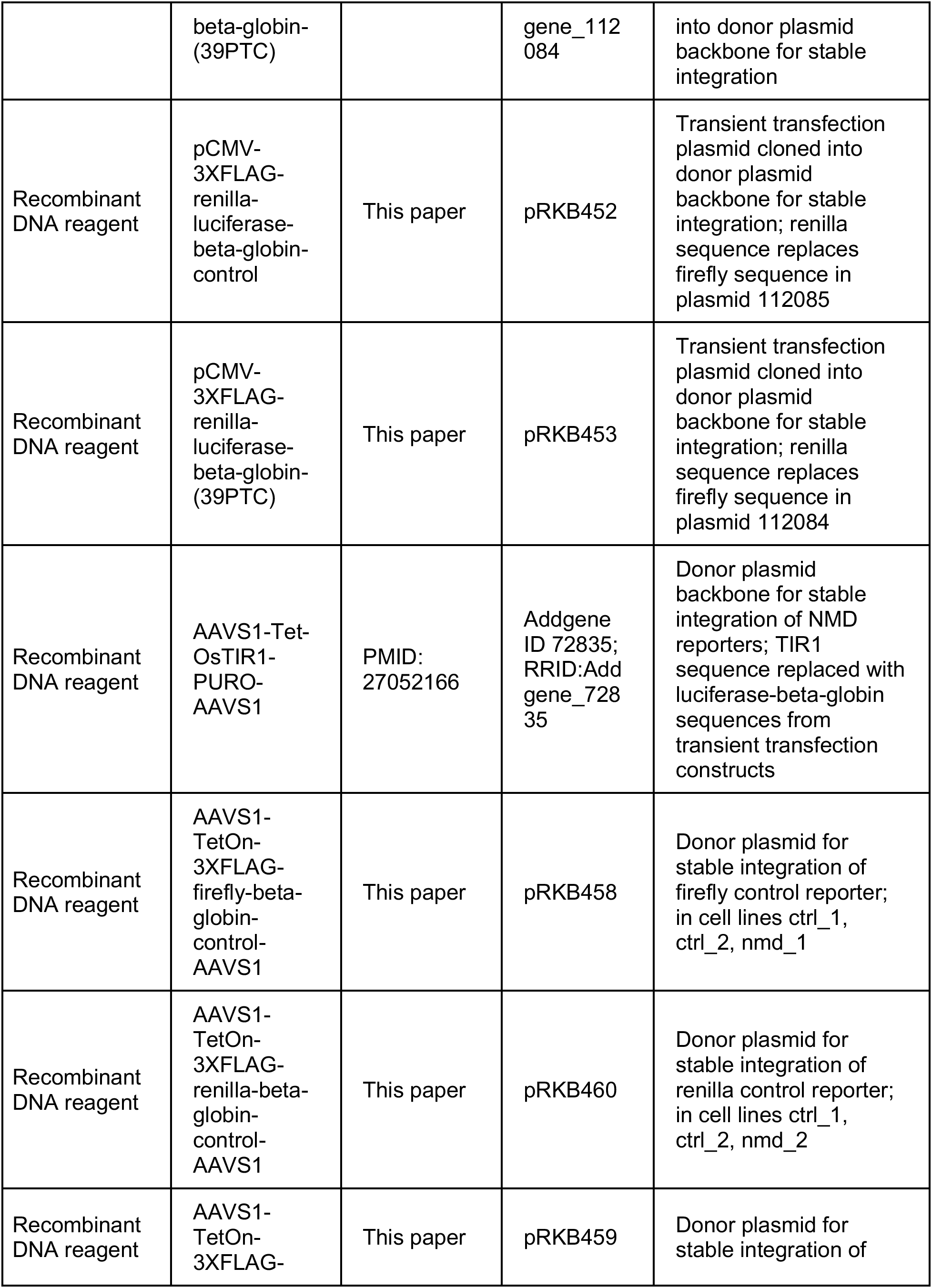
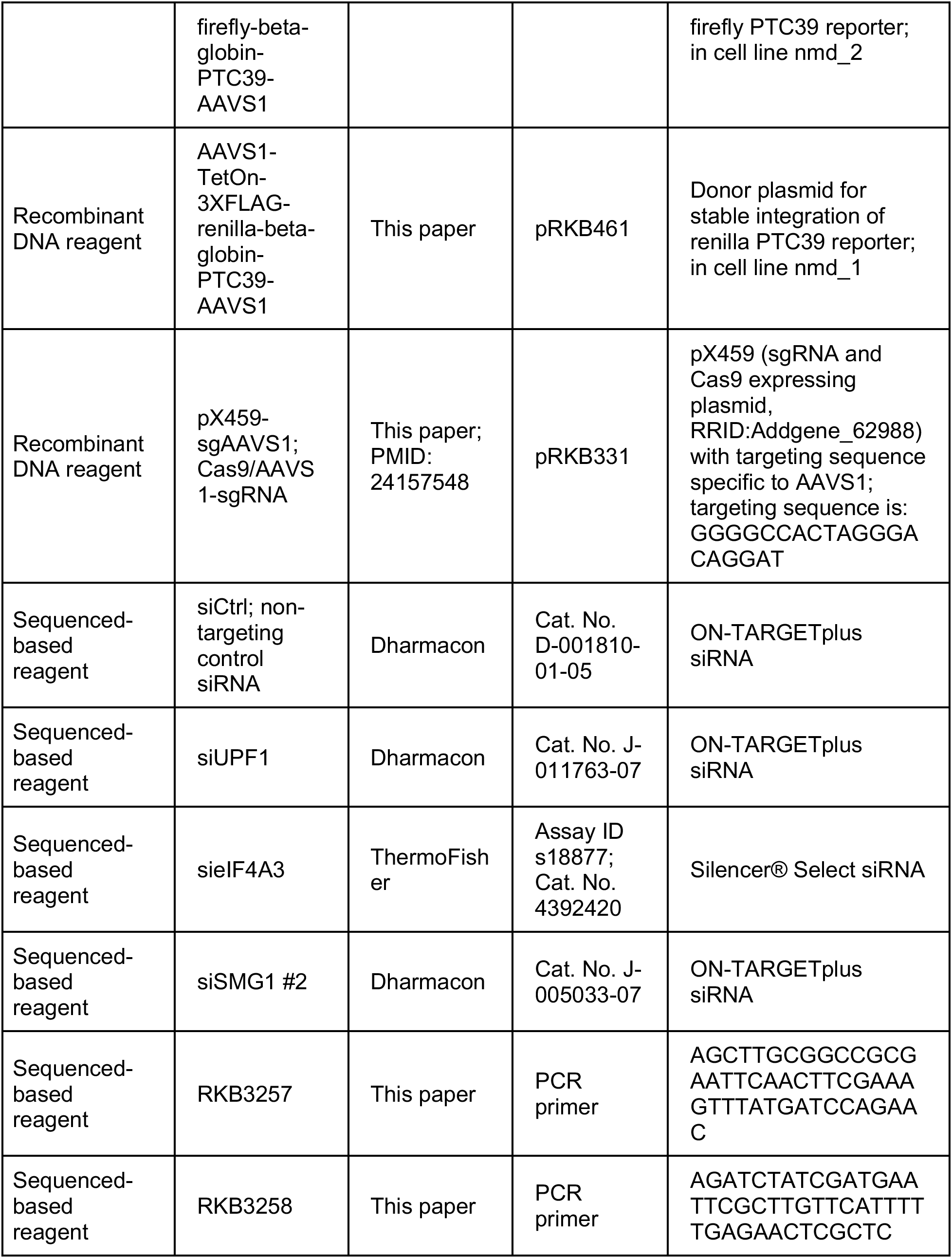
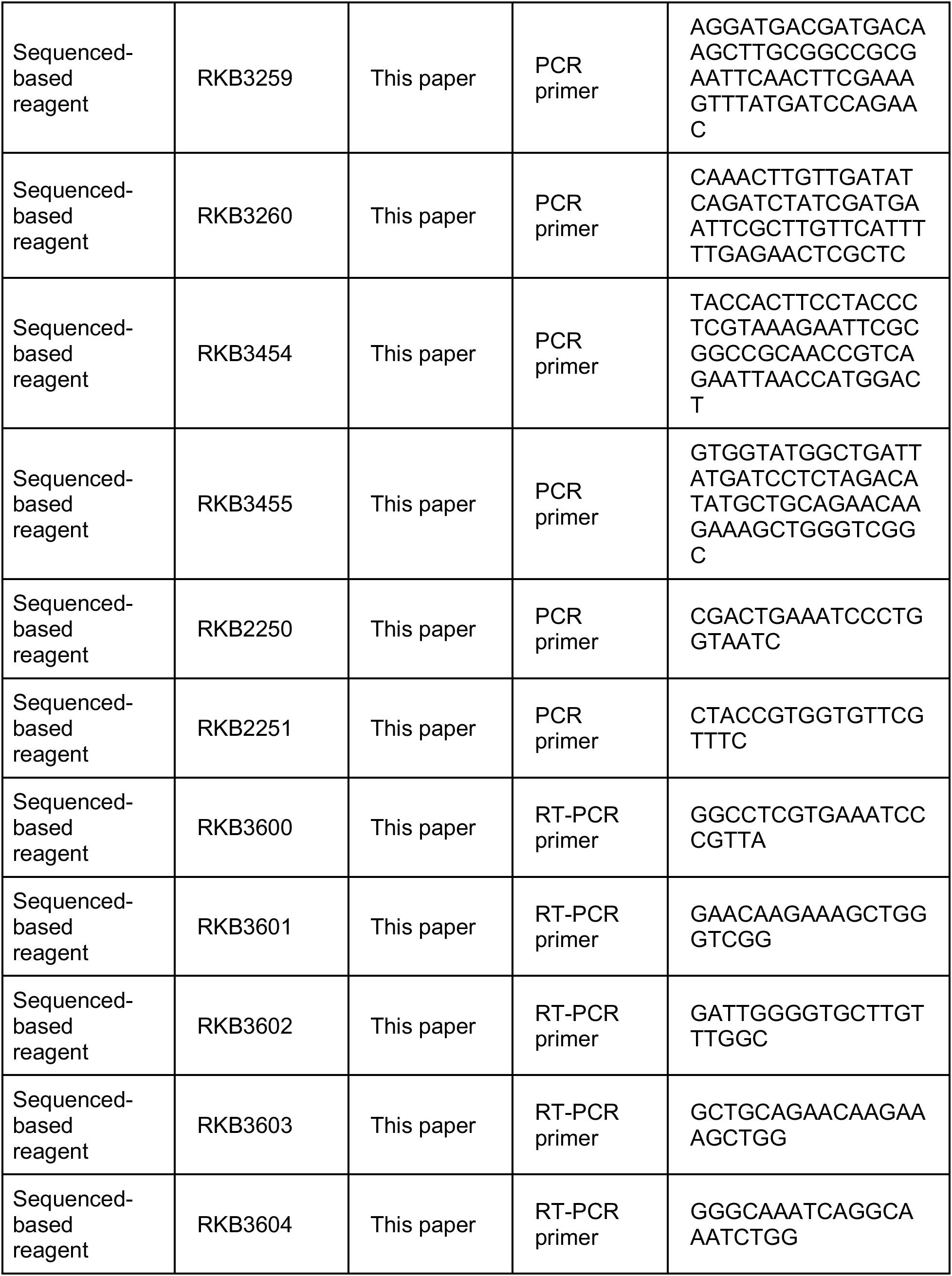
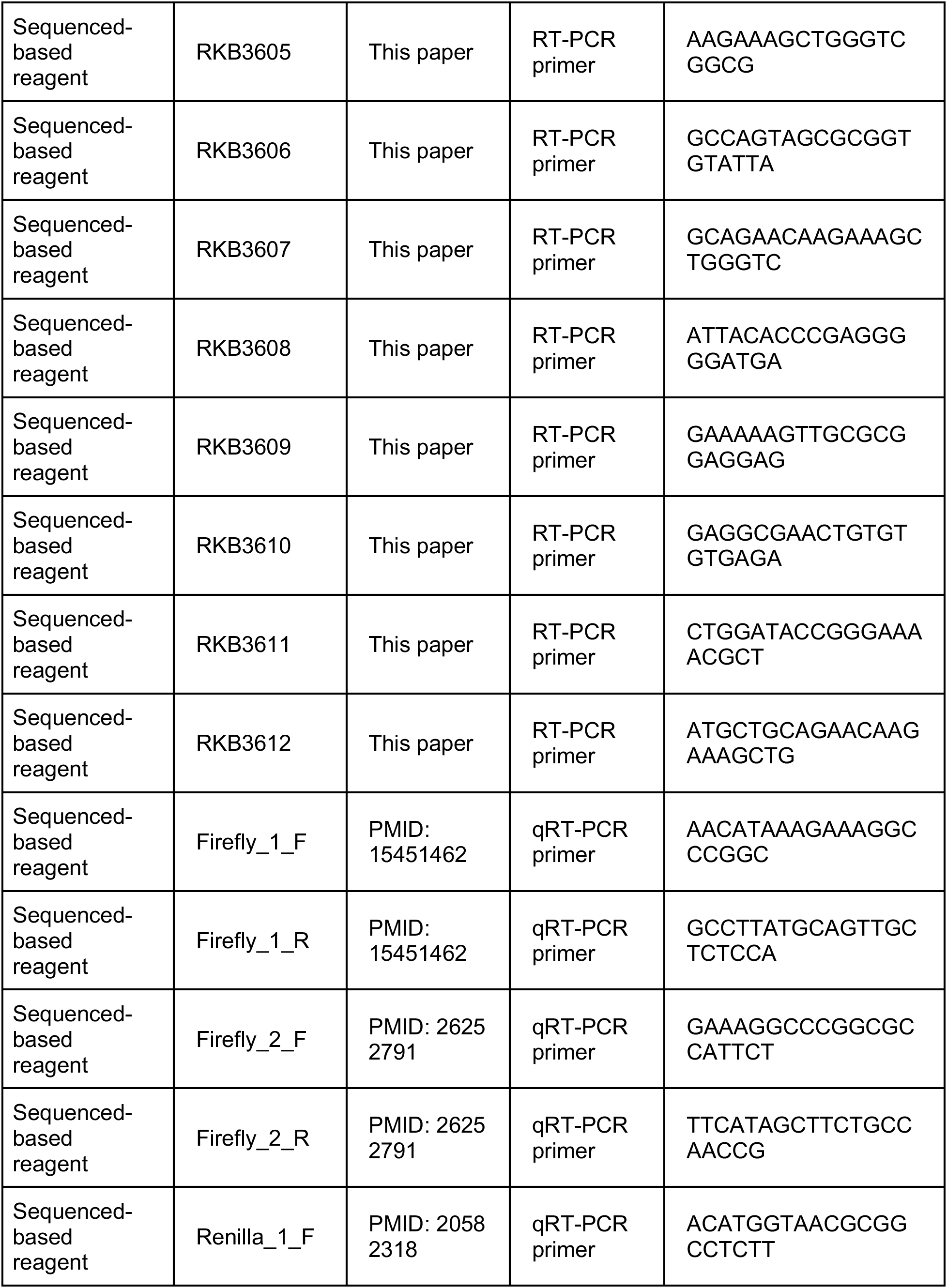
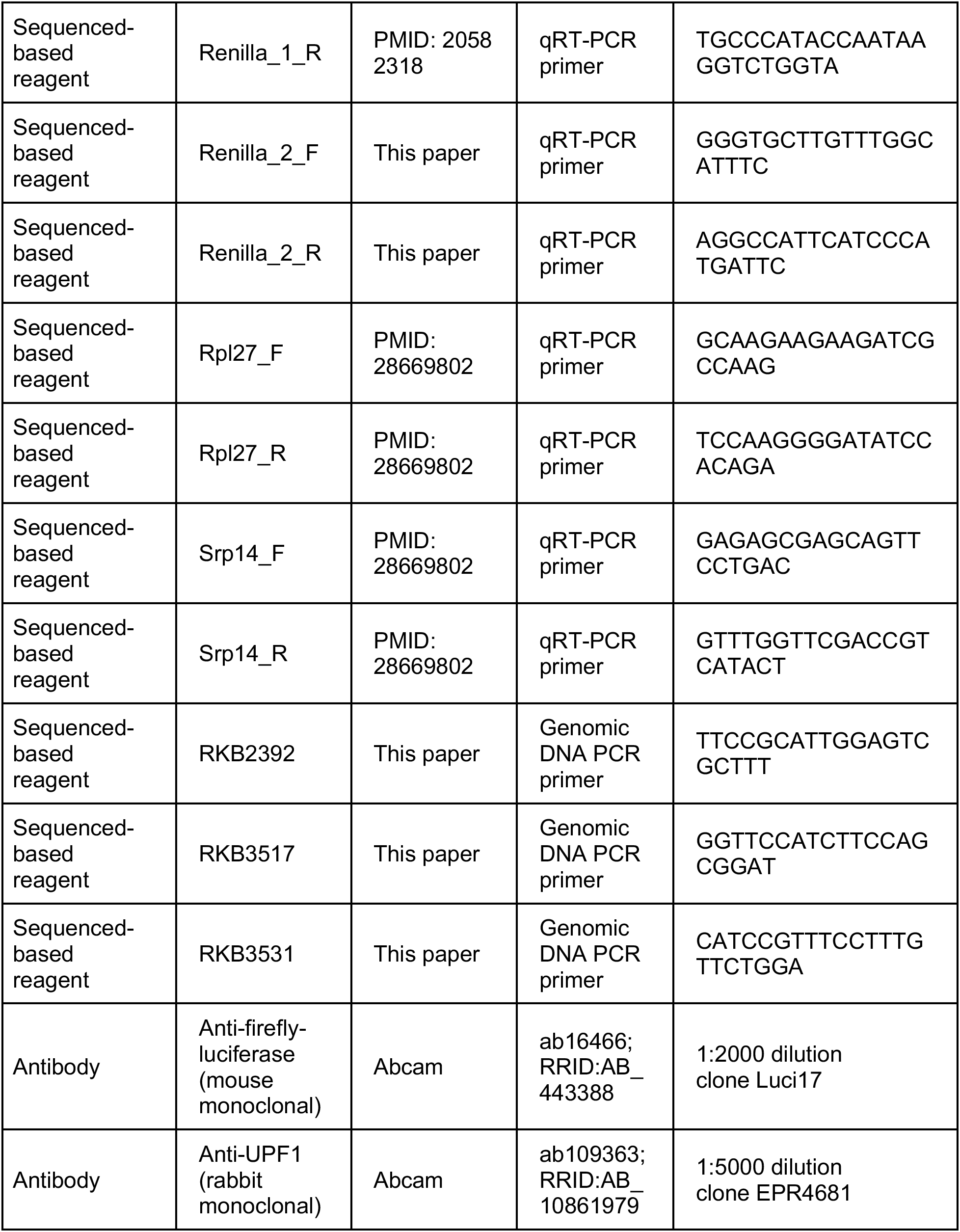
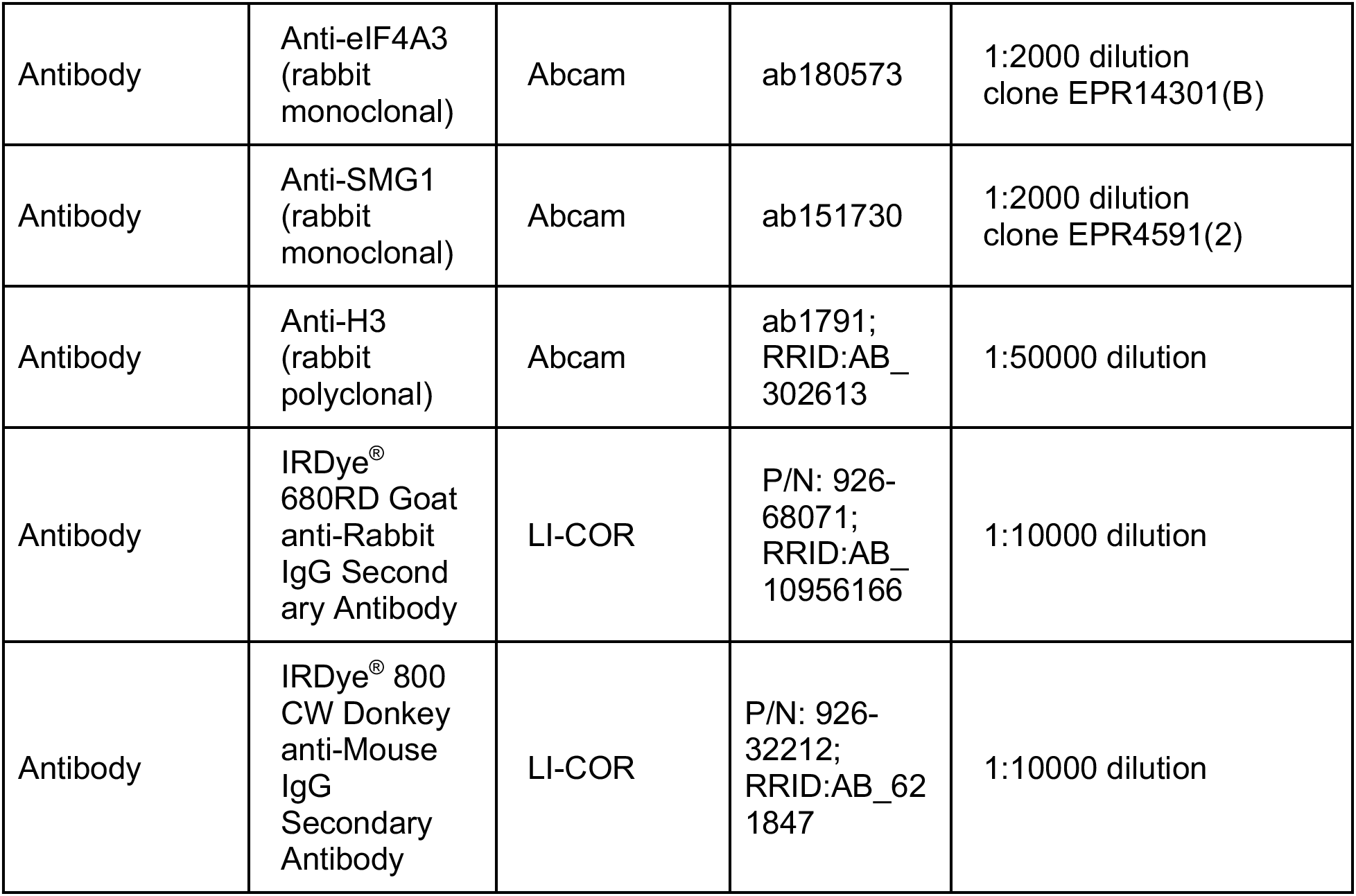

